# Sequence based mapping identifies *AWNS1*, a candidate transcription repressor underlying awn suppression at the *B1* locus in wheat

**DOI:** 10.1101/519264

**Authors:** Noah DeWitt, Mohammed Guedira, Edwin Lauer, Martin Sarinelli, Priyanka Tyagi, Daolin Fu, QunQun Hao, J. Paul Murphy, David Marshall, Alina Akhunova, Katie Jordan, Eduard Akhunov, Gina Brown-Guedira

**Affiliations:** Department of Crop and Soil Sciences, North Carolina State University, Raleigh, NC, USA 27695; Department of Plant Sciences, University of Idaho, Moscow, ID, USA 83844; USDA-ARS SAA, Plant Science Research, Raleigh, NC, USA, 27695; Department of Plant Pathology, Kansas State University, Manhattan, KS 66506

**Author notes:** Corresponding author: Gina Brown-Guedira, 01-919-513-0696.

## Abstract

Awns are stiff, hair-like structures that grow from the spikelets of wheat (*Triticum aestivum* L.) and other grasses. In wild wheat relatives, awns play a role in seed dispersal. Awn suppression in domesticated wheat, where awns are shortened or even eliminated entirely, is variably adaptive with both awned and awnless types under widespread cultivation. Although the *B1* locus on the long arm of chromosome 5A is a major determinant of awn suppression, no underlying gene or mechanism of action has been identified. Using association mapping, we identified a SNP marker located on the distal end of 5AL (5A28417) predictive of awn status in a panel of 640 U.S. winter wheat breeding lines, indicating that *B1* was the determinant of awn suppression in this germplasm. Analysis of historical data available for the panel determined *5A28417* was also significantly associated with grain test weight. Evaluation of spike morphology and kernel traits was undertaken in a 341 RIL population developed from a cross between awned soft winter wheat cultivar LA95135 and awnless cultivar SS-MPV57. Awn suppression in the population co-segregated with 5A28417 and was co-located with QTL for number of spikelets per spike, kernel weight and kernel length. Fine-mapping located *B1* to a region containing only two predicted genes, including a C2H2 zinc finger transcription factor 219 bp from 5A28417 that we named *AWNS-A1*. Deletions encompassing both genes were present in awned mutants of the awnless cultivar Brundage. Polymorphisms in the *AWNS-A1* coding region were not observed in diverse wheat germplasm. However, sequencing of wheat lines representing different marker haplotypes in the surrounding region identified a deletion 3 kb downstream of *AWNS-A1.* A marker for this deletion was highly predictive of awn suppression in a collection of diverse wheat accessions, and indicates that *AWNS-A1* is likely the major determinant of awn suppression in global wheat germplasm. *AWNS-A1* is more highly expressed in developing spikes of awnless individuals, suggesting a mechanism for awn suppression.

## Introduction

Wheat (*Triticum aestivum* L.) is a major food crop, producing roughly 20% of human calories and protein worldwide (FAOSTAT 2018). Given the dual challenges of human population growth and the negative impacts of climate change, increasing yield of this staple food crop is essential to ensuring food security. A key factor in wheat’s historic spread and current global importance is its ability to adapt to a range of environments; however, wheat grows best in cooler climates, and exposure to higher temperatures during grain fill are detrimental to grain yield and quality. As temperatures rise globally, identifying and exploiting the genetic basis for yield has become a major focus of wheat breeding. Genetic control of grain yield is highly polygenic. One approach to exploring the genetics of this complex trait is to dissect grain yield into simpler phenotypes and study the genetic architecture behind each of these yield components. Key components of grain yield in wheat include grain number per unit area and, to a lesser extent, grain size (Kuchel *et al.*, 2007).

Awns are stiff, hair-like structures common in grass inflorescences. In *Poaceae*, awns emerge from the lemma of young spikelets at an early developmental stage. Awns play an important role in seed dispersal in wild relatives of wheat with a brittle rachis; seeds attach to passing animals via small, stiff hooks lining the awns. In addition, awns balance seeds as they fall, and their expansion and contraction in response to changes in humidity drives seeds into the soil (Elbaum *et al.*, 2007). Awns may also deter herbivores from ingesting seed heads (Grundbacher 1963), making awn suppression important for developing forage cultivars (Cash *et al.*, 2009). The absence of awns in rice is considered a key domestication trait facilitating grain harvest and storage (Toriba & Hirano 2013), but the history of the evolution and spread of awn suppression in wheat is not well understood.

Wild-type wheat is awned, with major variation for awn length emerging from different combinations of three dominant genes: *B1 (Tipped 1)*, *B2 (Tipped 2)* and *Hd (Hooded)* (McIntosh *et al.*, 2013). On its own, the *B1* awn suppressor produces an apically awnletted phenotype, characterized by short awns at the end of the spike and absent elsewhere (Watkins and Ellerton, 1940). The *B2* allele reduces awn length most dramatically towards the top and bottom of the wheat spike, while the *Hd* allele reduces awn length consistently and can produce curved, “hooked” awns (Watkins and Ellerton, 1940). A combination of *Hd* and *B1* or *Hd* and *B2* (as in the model wheat cultivar Chinese Spring) is required for a mostly awnless phenotype, and a combination of all three alleles is needed for complete awn suppression (Yoshioka *et al.*, 2017). None of the genes controlling awn length in wheat have been cloned, but mapping studies place *Hd* and *B2* on the short arm of chromosome 4A and the long arm of chromosome 6B, respectively (Sourdille *et al.*, 2002, Yoshioka *et al.*, 2017). The *B1* locus has long been used as a physical marker, located distal to the major genes controlling vernalization requirement (*VRN-A1)* and the spelt head type (*Q*) on 5AL (Kato *et al.*, 1998). Recent fine mapping has narrowed the *B1* region to a 7.5 cM interval on the distal end of 5AL closely linked to marker *BW8226_227* (Mackay *et al.*, 2014). In *Aegilops tauschii*, the donor of the D-genome in hexaploid wheat, an additional dominant awn suppressor *Anathera* (*Antr*) was located on the distal end of 5DS (Nishijima *et al.*, 2017). Deletion of the short arm of chromosome 3B in Chinese Spring also produces an awned phenotype, suggesting that further uncharacterized genes are involved in awn development (Ma *et al.*, 2012).

Awn suppression in barley is controlled by a *Hooded* gene (*K*) that replaces awns with sterile flowers, and a *short awn 2* gene (*lks2*) which shortens awns (Roig *et al.*, 2004, Takahashi 1955). These genes have been cloned and identified as genes involved in the awn developmental pathway: a homeobox gene *Knox3* overexpressed due to an intron duplication (Müller *et al.*, 1995) and a *SHORT INTERNODES* transcription factor (Yuo *et al.*, 2012). Awn suppression is an important domestication trait in rice, and has been shown to also result from mutations in genes involved in awn development (Luo *et al.*, 2013, Toriba & Horina 2013). However, awn supppression in wheat does not appear to be related to these developmental genes (Yoshioka *et al.*, 2017).

Variation in awn length across modern wheat cultivars and landraces, including within both spring and winter wheat, suggests awns are variably adaptive in different environments. Previous studies support a role for awns in supplying photosynthate to developing seeds of wheat and barley (Grundbacher 1963, Kjack & Witters, 1974, Motzo & Giunta, 2002, Li *et al.*, 2006, Tambussi *et al.*, 2007, Ali *et al.*, 2010, Maydup *et al.*, 2014), and the location of awns on the wheat head facilitates movement of carbohydrates into seeds and positions them favorably for photosynthesis (Ali *et al.*, 2010, Evans *et al.*, 1972, Li *et al.*, 2006). Awns may continue contributing to photosynthesis if leaves senesce early or are damaged by disease or drought (Tambussi *et al.*, 2007), and the absence of awns can halve the rate of net ear photosynthesis (Evans *et al.*, 1972). Besides increased area for photosynthesis, in warmer climates awns can play a role in cooling the wheat spike during grain fill (Motzo & Giunta, 2002). Awned wheats have been demonstrated to perform better in hotter climates or under water stress. The presence of awns is associated with smaller numbers of larger seeds (Rebetzke *et al.*, 2016). Awn tissue, potentially due to its silica coating, tolerates water deficit better than other important photosynthetic tissues such as the flag leaf (Tambussi *et al.*, 2005, Peleg *et al.*, 2010).

However, awnless or awnletted wheat types are widely cultivated and are the dominant morphology in many parts of the world. The absence of awns is associated with a reduced incidence of pre-harvest sprouting (Cao *et al.*, 2016, King & Richards, 1984). Other potential explanations for the prevalence of awnless varieties include use of wheat as forage and historical ease of harvest. In warm growing regions like the southeastern U.S. where awnless varieties are historically dominant, the proportion of awned varieties has increased over the past two decades. Given the potential influence of awns on development of the wheat spikelet and kernels, a better understanding of the genetic basis of awn suppression should facilitate an understanding these processes. With the proliferation of methods developed alongside next generation sequencing technology and the publishing of a reference genome, the wheat genetics community has developed a number of freely available genomics resources (Appels *et al.*, 2018). The objective of this study was to utilize these new resources to identify a candidate gene responsible for awn suppression at the *B1* locus.

## Materials and Methods

### Genome Wide Association Analyses

An association mapping panel was evaluated that consisted of 640 elite soft winter wheat breeding lines. These lines were entries in two collaborative testing nurseries in the southeast soft wheat growing region of the United States, the Gulf Atlantic Wheat Nursery (GAWN) and the Sunwheat Nursery, over a period of nine growing seasons (2008 to 2016). Two replications of each line were grown in 1-meter rows spaced 30 cm apart at Raleigh, North Carolina during the 2016-2017 growing season. At heading, presence or absence of awns was noted.

Association analyses were performed for test weight using historical data available for the 640 entries. The GAWN and SunWheat nurseries were evaluated at one location in up to seven states per year from 2008 to 2016: Arkansas (Stuttgart or Marianna), Florida (Citra or Quincy), Georgia (Plains), Louisiana (Winnsboro), North Carolina (Kinston), Texas (Farmersville) and Virginia (Warsaw). Experimental designs at each environment were randomized complete block designs with one to three replications. Plot size was typical of yield trial plots for wheat in the region with a minimum of 1.3 meters wide and 3.1 meters long. The data set was balanced for individual years where the same set of genotypes was evaluated across different locations and unbalanced between different years.

The following linear mixed model was utilized for the analysis of test weight:

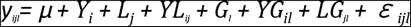

Where, y_*ijl*_ was the phenotypic observation of genotype *l* in the *i*^*th*^ year in the *j^th^* location, *μ* was the overall mean, Y_*i*_ was the year effect, L_*j*_ was the location effect, G_*l*_ was the genotypic effect, YL_*ij*_, YG_*il*_, LG_*jl*_, were the interaction terms representing year by location, genotype by year, genotype by location, respectively and *ε_ijl_*, represented the residual term. For this model, the overall mean and the genotypic effect were considered fixed and all the remaining terms random. A homogeneous residual error variance term was fitted: ε~IID 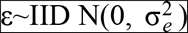 due to the lack of replications within each environment. Random effects (u) and residuals (e) were assumed to be normally and independently distributed û~IID 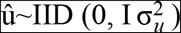 The models were implemented using ASReml-R (Butler *et al*., 2009). Best linear unbiased estimates (BLUE) of each genotype were calculated as the estimated genotypic effect plus overall mean and used as the response variable for association mapping.

Seed of each entry was germinated and DNA was extracted using the sbeadex plant maxi kit (LGC Genomics LLC, Teddington, UK). Genotyping by sequencing (GBS; Elshire *et al.*, 2011) was performed according to Poland et al. (2012), with ninety-six individual samples barcoded, pooled into a single library, and sequenced on an Illumina HiSeq 2500. Tassel5GBSv2 pipeline version 5.2.35 (Glaubitz *et al.*, 2014) was used to align raw reads to the International Wheat Genome Sequencing Consortium (IWGSC) RefSeqv1.0 assembly (https://wheat-urgi.versailles.inra.fr/Seq-Repository/Assemblies) using Burrows-Wheeler aligner (BWA) version 0.7.12 and call SNPs (Li *et al.*, 2009). SNPs were filtered to retain samples with ≤ 20 percent missing data, ≥ 5 percent minor allele frequency and ≤ 10 percent of heterozygous calls per marker. Missing SNPs were imputed using Beagle (Browning & Browning, 2016).

GWAS for awn status and test weight were conducted in R version 3.3.1 (R Core Team 2016) using the package genome association and prediction integrated tool GAPIT (Lipka *et al.*, 2012). Manhattan plots were produced in R using the package qqman (Turner 2014). Population structure and degree of relatedness between individuals were accounted for using the first three principal components and the genomic relationship matrix determined using the GBS markers. Principal Components Analysis of this matrix was implemented with the prcomp function in R version 3.3.1 (R Core Team 2016). Markers were declared significant based on the Bonferroni corrected p-value at alpha = 0.01.

### Bi-Parental Mapping Populations

A population of 341 F_5_-derived RILs was developed from the cross of awned cultivar LA95135 with awnless SS-MVP57 (LM population). LA95135 possesses the *Rht-B1a* (wild type) and *Rht-D1b* (dwarfing) alleles and the *Ppd-A1a.1* (Nishida *et al.*, 2013) allele for photoperiod insensitivity. SS-MPV57 had wild type alleles *Rht-B1a* and *Rht-D1a* (Guedira *et al.*, 2010) and the *Ppd-D1a* allele conferring photoperiod insensitivity.

During the winter of 2016-2017, an experiment was conducted in the greenhouse to evaluate spike morphology and kernel weight. Imbibed seeds from each RIL were placed in a cold chamber kept at 4°C for 8 weeks and were transplanted into plastic cones (Volume 0.7L, 6.5 cm in diameter and 25 cm depth) containing soil mix. Plants were grown in a completely randomized design having four replications in a greenhouse set at 16 hr photoperiod and 20°C/15°C (day/night) temperature. The main tiller of each plant was used for evaluation of spike length, spikelet number per spike, kernels per spike, kernel weight and presence or absence of awns.

The LM RIL population was evaluated in the field at Raleigh, NC and Kinston, NC during the 2017-2018 season. The 341 RILs were grown in an incomplete block experiment with two replications at each location. The population was divided into five blocks, each consisting of 72 entries and the two parents of the population. Plots consisted of 1-meter rows spaced 30 cm apart. Six representative spikes from each row were harvested and number of spikelets recorded. Seeds were weighed and counted to determine kernel weight. Rows at the Raleigh, NC location were hand harvested and threshed using a Vogel thresher. A 15 ml sample of seed was cleaned and grain morphometric parameters (grain length, width, area and weight) were obtained using a MARVIN grain analyzer (GAT Sensorik GMBH, Neubrandenburg, Germany). An estimate of test weight was obtained by measuring the weight of seed in the 15 ml sample. BLUEs were calculated for individual RILs using the R package lme4, treating genotype as a fixed effect and all other terms as random (Bates *et al.*, 2015).

RILs and parents of the LM population were genotyped and SNP identified using the GBS protocol as described above, except that missing data were not imputed. A linkage map was constructed with these data, KASP markers for major effect loci *Rht-D1* and *Ppd-D1*, and the presence or absence of awns as a physical marker using the R packages R/qtl and ASMap (Broman *et al.*, 2003, Taylor *et al.*, 2017). QTL analysis was performed using composite interval mapping, with significance thresholds for an alpha = 0.05 determined using 1000 permutations.

Three F2 populations (referred to as GxM, GxS, and LxS) were developed from crosses between awned breeding lines GA06493-13LE6 (G) and LA09264C-P2 (L) with the awnless cultivars SSMVP-57 (M) and SS8641 (S). Imbibed F2 seed were vernalized for 6 weeks, transplanted into plastic cones and grown in a greenhouse as described above. DNA was isolated from seedling tissue using the sbeadex plant maxi kit (LGC Genomics LLC, Teddington, UK). After heading, a total of 950 plants were evaluated for presence or absence of awns.

Genomic DNA of the RIL and F2 populations was evaluated with KASP markers located in the *B1* region. Markers were developed from sequences flanking GBS SNPs in the *B1* region (Tables S1 and S2) in the LM mapping population and SNPs identified in the exome capture database developed for the WheatCAP project that included LA95135 and SS-MPV57 (https://triticeaetoolbox.org/wheat/display_genotype.php?trial_code=2017_WheatCAP). A linkage map of the distal region of the long arm of chromosome 5A containing *B1* and selected KASP assays for each F_2_ and the RIL population was developed using R/qtl (Broman *et al.*, 2003).

Scaffold assemblies of awnless winter wheats with the *B1* suppressor (Cadenza, Paragon, Robigus, and Claire) and the awned tetraploid wheat Kronos were downloaded from the Earlham Institute website (https://opendata.earlham.ac.uk/opendata/data/). Scaffolds containing portions of the B1 region in the awnless assemblies and Kronos were identified using BLAST and aligned to the IWSGC reference assembly RefSeqv1.0 of Chinese Spring using LAST (Kielbasa *et al.*, 2011). Samtools was used to identify SNPs between the awned and awnless wheats for KASP marker design (Li *et al.*, 2009).

### Analysis of mutant lines

Seeds of the awnless cultivar Brundage were treated using fast neutron irradiation performed at the McClellan Nuclear Radiation Center (McClellan, CA, USA) with a center total dose of 7 Gy air. M1 seeds were planted in the Parker Farm (Moscow, ID, USA) during the 2016-2017 crop season. The main tiller was harvested from each M1 plant and seeds were planted as one meter rows at the Parker Farm in the 2017-2018 crop season. At heading, rows were noted as being either awnless or segregating for awns. Markers in the B1 region were evaluated on DNA isolated from one awnless plant and up to four awned plants from segregating rows. DNA samples of Chinese Spring and the Chinese Spring deletion line 5AL-6 (TA4535-6) missing the terminal 32% of 5AL were included as controls to evaluate genome specificity of markers. Progenies from the family with the smallest deletion around *B1* were grown in the greenhouse in Raleigh in 2018 along with Chinese Spring and TA4535-6 to assess the awn phenotype. KASP assays for SNP located on chromosomes 3B, 1A, 4A, and 6A were evaluated on all mutants to exclude the possibility of off-type plants in the field.

### Identification of haplotypes in diverse wheat germplasm

Genomic DNA from a panel of 455 winter and 1984 spring wheat accessions representing global diversity as part of the core collection of the USDA-ARS National Small Grains Collection (NSGC) was evaluated using KASP markers developed from a 127 kb region flanking the *B1* locus. Data on the presence or absence of awns for winter wheat accessions was gathered from the U.S. National Plant Germplasm System website (https://npgsweb.ars-grin.gov). The spring wheat accessions were grown as single 1-meter rows spaced 30 cm apart at Raleigh, NC and were planted during October 2017. At heading time, single tillers were selected for each accession, the presence or absence of awns was noted and genomic DNA was isolated from the flag leaf.

### Sanger Sequencing *AWNS1*

The two parents of the RIL population and eight individuals representing diverse haplotypes for markers flanking *B1* were selected for Sanger sequencing of the *AWNS1* candidate gene (Table S3). Amplification of the coding sequence and 7 kb of surrounding non-repetitive sequence was performed with a nested PCR design to obtain specificity using NEB Longamp polymerase (New England Biolabs, Ipswich, MA). The region was divided into 2.5 kb and 4.5 kb regions, and two forward and reverse primers were developed for each region. PCR products were run on a 2% agarose gel with ethidium bromide to gauge specificity and fragment length. PCR using the outer set of forward and reverse primers produced multiple PCR products, including fragments of the target size. Purified PCR products were used as a template for amplification with the inner set of forward and reverse primers, producing a single band of the expected size. Forward sequencing primers were designed every 500-800 bp for Sanger sequencing of the region. CodonCode Aligner software (CodonCode Corporation, www.codoncode.com) was used to check base calls and assemble sequence reads into contigs.

### Gene Expression

To evaluate expression of candidate genes, tissue from immature inflorescences was collected from primary and secondary tillers of plants of LA 95135 and SS-MPV57, as well as the awned cultivar AGS 2000 and awnless cultivar Massey. Individual samples from spikelets at similar stages were grouped for analysis to assess developmental variation in gene expression. RNA was isolated from plant tissue using the Zymopure RNA Extraction kit (Zymo Research, Irvine, CA), and reverse transcribed with the ThermoFischer Reverse Transcription kit. The predicted exon of candidate gene TraesCS5A02G542800 was aligned to orthologous sequences on chromosomes 4B (not annotated) and 4D (TraesCS4D01G476700LC) to design genome-specific primers for qPCR with an amplicon size of 100-150 bp and a T_m_ of 60-62° C. Non-genome specific primers for *β-ACTIN* were selected as an endogenous control (Li *et al.*, 2013). PCR was performed using the designed primers (Table S4) on converted cDNA using NEB taq polymerase (New England Biolabs, Ipswich, MA), and run on a 2% agarose gel with ethidium bromide to test for specificity.

Quanitative PCR reactions were performed using a CFX384 real time PCR machine (Bio-Rad Laboratories, Hercules, CA) with Sybr green qPCR Master Mix (Applied Biosystems, Foster City, CA). Reactions included primers for the candidate gene along with a *β-ACTIN* control at an annealing temperature of 61° C. Three technical replications were performed per sample. Cq values were calculated for each replication using the Biorad CFX Maestro software and normalized to expression relative to the endogenous control *β-ACTIN* with 2^^(ACTIN CT - TARGET CT)^, representing the number of molecules of the target gene compared to *β-ACTIN*.

As an additional evaluation of gene expression over the course of apical development, the sequences of genes in the B1 region were submitted to the WheatExp wheat expression database (https://wheat.pw.usda.gov/WheatExp/). *β-ACTIN* primer sequences were also submitted to verify that its expression is consistent during apical development.

## Results

### Predictive SNP for *B1* awn supression is associated with test weight, spikelets per spike and kernel weight

Of the 640 lines evaluated in the eastern soft winter wheat panel, 58 % had awns. Association mapping utilizing 14,567 GBS markers identified 30 markers significantly associated (adjusted P-value < 0.01) with the presence or absence of awns (Table S5). The significant markers were in a distal region of the long arm of chromosome 5A, consistent with the location of the *B1* awn suppressor (Fig. **1a**). Alignment of markers to IWGSC RefSeqv1.0 of Chinese Spring wheat placed the *B1* locus in a 25 Mb region between 681,455,268 bp and 706,705,101 bp. A SNP located at 698,528,417 bp on chromosome 5A (5A28417) was highly significant (P-value = 7.19 x 10^−57^) and 100% predictive of awn status in this set of lines. Association analysis of historical data for test weight of the GAWN and SunWheat regional testing nurseries over the period from 2008 to 2017 identified a significant association of test weight with SNP 5A28417 (P-value = 4.14 x 10^−7^; Fig. **1b**).

**Figure 1.**
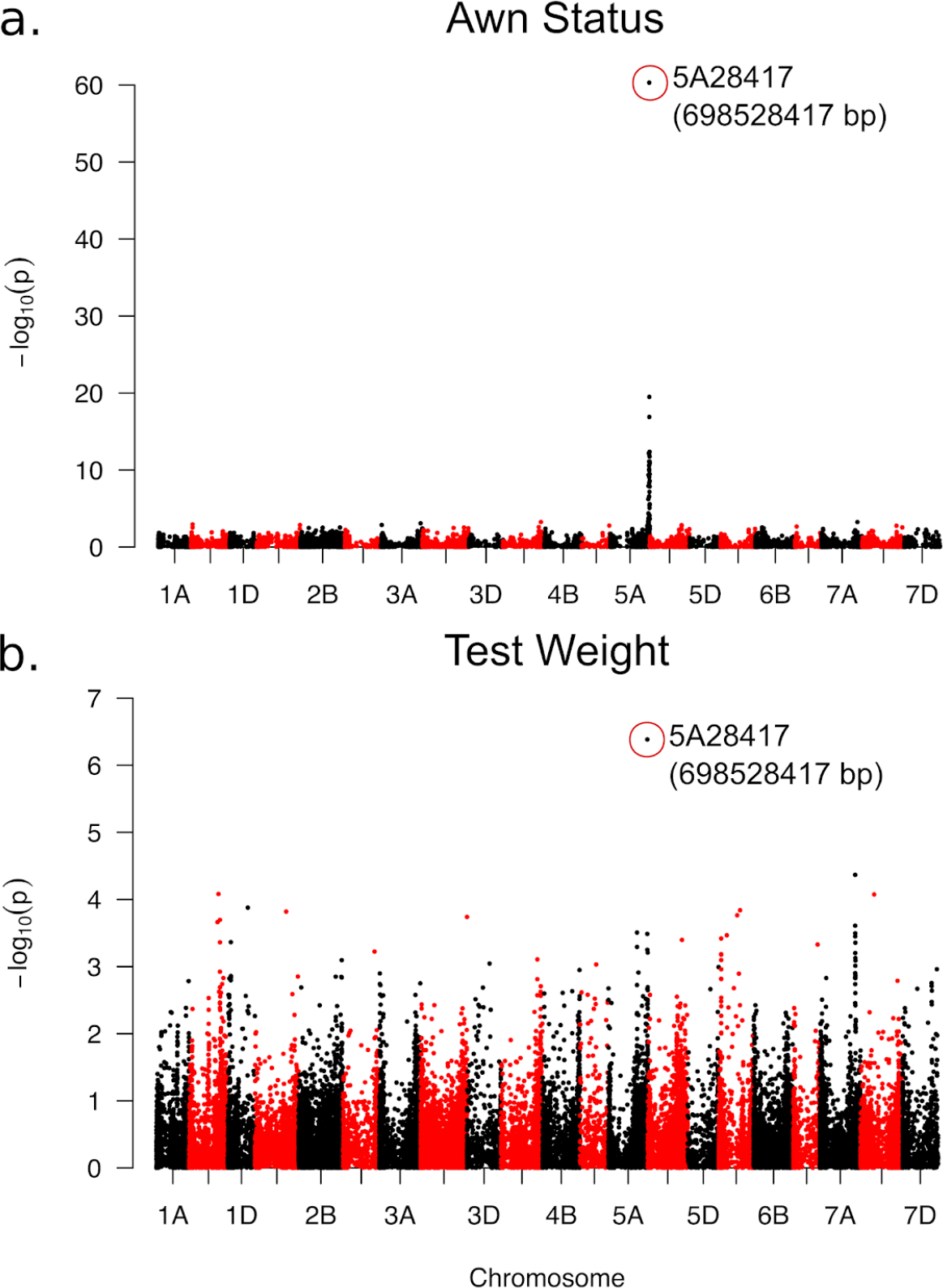
GWAS analysis conducted for awn status (a) and test weight (b). The most significant marker awn status (*5A28417*) is also the most predictive marker for test weight.

Analysis of the LM RIL population identified multiple QTL for spikelets per spike and kernel morphometric traits in the field and greenhouse experiments, including highly significant QTL associated the *B1* locus and with markers predictive of major effect photoperiod and plant height genes *Ppd-D1* and *Rht-D1* (Fig. 2, Table S6). The *Rht-D1b* semi-dwarfing allele was associated with 9.2% and 12.3% decreases kernel weight in the field and greenhouse, respectively, and associated with QTL for reduced kernel width and kernel area (Table S6). The *Ppd-D1a* allele for photoperiod insensitivity was associated with a 6.9% decrease in thousand kernel weight in the greenhouse and 2% and 6.1% decrease spikelets per spike in the field and greenhouse, respectively. The presence or absence of awns was significantly associated with spike length, spikelets per spike, kernel length and thousand kernel weight. In the greenhouse experiment, the presence of awns increased thousand kernel weight 1.47 g (5.1%), and decreased spikelets per spike by 0.48 spikelets (−2.7%). In the field experiment, the presence of awns increased thousand kernel weight by 0.88 g (3.2%), and decreased spikelets per spike by 0.37 spikelets (−1.8%). In addition, *B1* significantly associated with increases in estimated test weight and kernel length in 2018 field data (Fig. 2, Table S6). We did not observe a significant effect of the *B1* locus on kernel width or area (Table S6).

**Figure 2.**
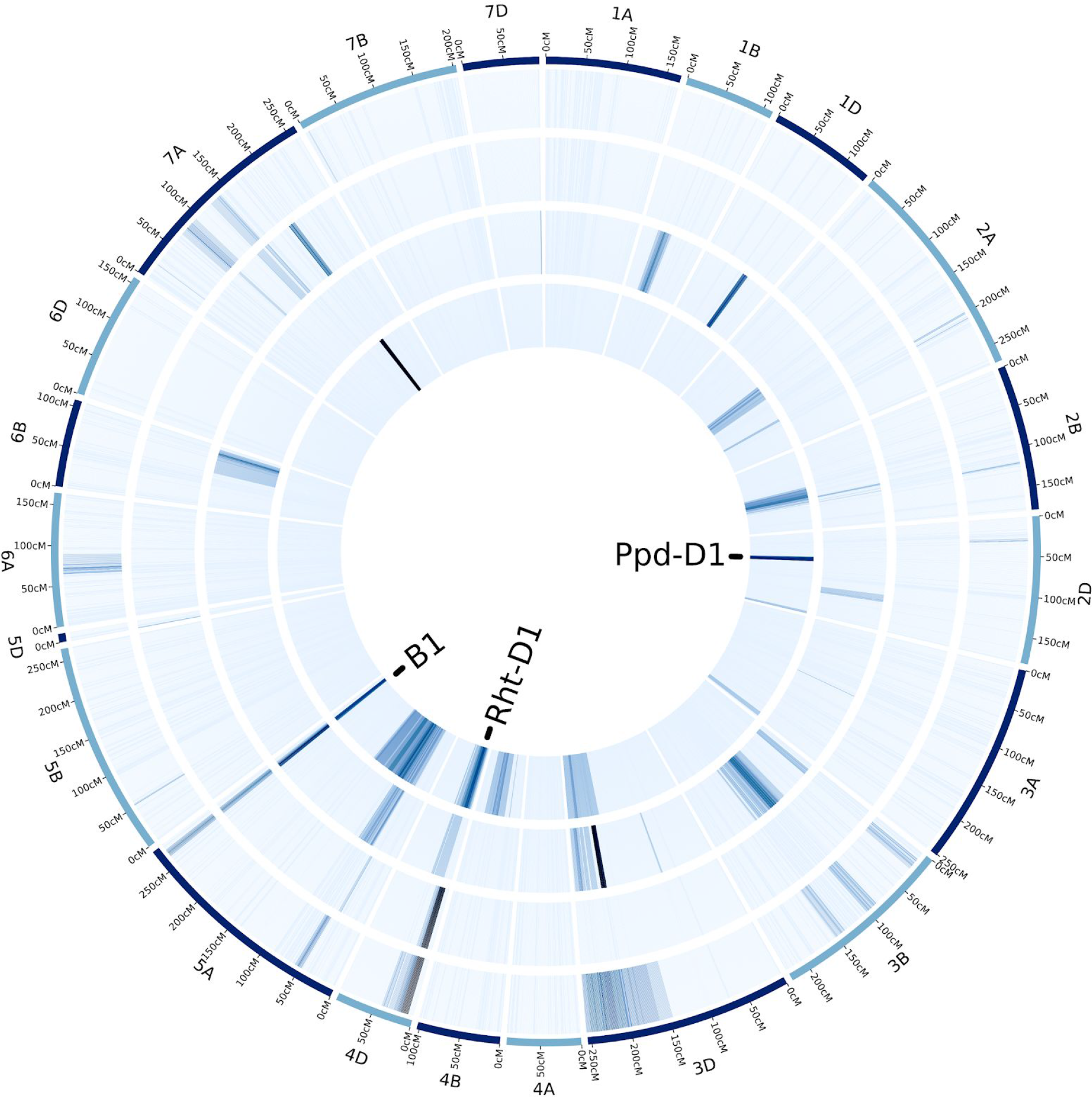
Heatmap of LOD scores for composite interval mapping in the LA95135 x SS-MPV57 RIL population of traits significantly impacted by presence or absence of awns. Traits from the outer to inner circle are thousand kernel weight, test weight, kernel length, and spikelets per spike. Location of the *Rht-D1*, *Ppd-D1* and *B1* loci are noted in center. Data on thousand kernel weight, test weight, and kernel length were collected from the field in Raleigh, NC in 2018. Data on spikelets per spike were collected from the field in Raleigh and Kinston, NC in 2018.

### Fine mapping identifies*B1* candidate genes

Significant SNP 5A28417 identified in the *B1* QTL region was located 219 bp upstream of predicted gene TraesCS5A02G542800 (Chr5A:698528636..698529001) in Chinese Spring RefSeqv1.0. The LM RIL population and three F_2_ populations developed from crosses between selected awned and awnless individuals in the association mapping panel were genotyped by KASP for SNPs targeted to an 8.52 Mb region flanking TraesCS5A02G542800 (Table S2). In each population, an awnless phenotype was correlated with a single gene co-segregating with KASP marker *5A28417*. Exome capture of the LA95135 and SS-MPV57 parents of the LM RIL population did not reveal polymorphisms in the TraesCS5A02G542800 coding sequence. Polymorphisms identified in predicted genes proximal (TraesCS5A02G542600 and TraesCS5A02G5426700) and distal (TraesCS5A02G542900) to TraesCS5A02G542800 were targeted for marker development. KASP marker *5A15019* targeted an A/G variant in intron five of predicted gene TraesCS5A02G542700 and *BW8226_227* targeted a T/G polymorphism in exon two of TraesCS5A02G542600. Marker *5A30334* targeted a C/T variant in exon one of predicted gene TraesCS5A02G542900.

Recombination within the bi-parental populations narrowed the genomic region to a 127 kb region containing two predicted genes (Fig. 3). Recombination events were observed between the awn phenotype and marker *5A30334* in the LM RIL population, locating the SNP in TraesCS5A02G542900 0.2 cM distal to *B1*. No recombination was observed between *B1* and *5A28417*, *5A15019* and *BW8226_227* in the LM population. Of the 950 individuals evaluated from the three F_2_ populations, individual GM#101 derived from the cross between GA06493-13LE6 and SS-MPV57 was determined to be homozygous for *BW8226_227* and heterozygous for *5A28417* and *5A15019*. The awn phenotype segregated in a progeny test of 16 F_3_ plants derived from GM#101, placing the SNP in TraesCS5A02G542600 0.2 cM proximal to *B1*. These results narrowed candidate genes underlying *B1* awn suppression to predicted genes TraesCS5A02G542700 and TraesCS5A02G542800.

**Fig. 3.**
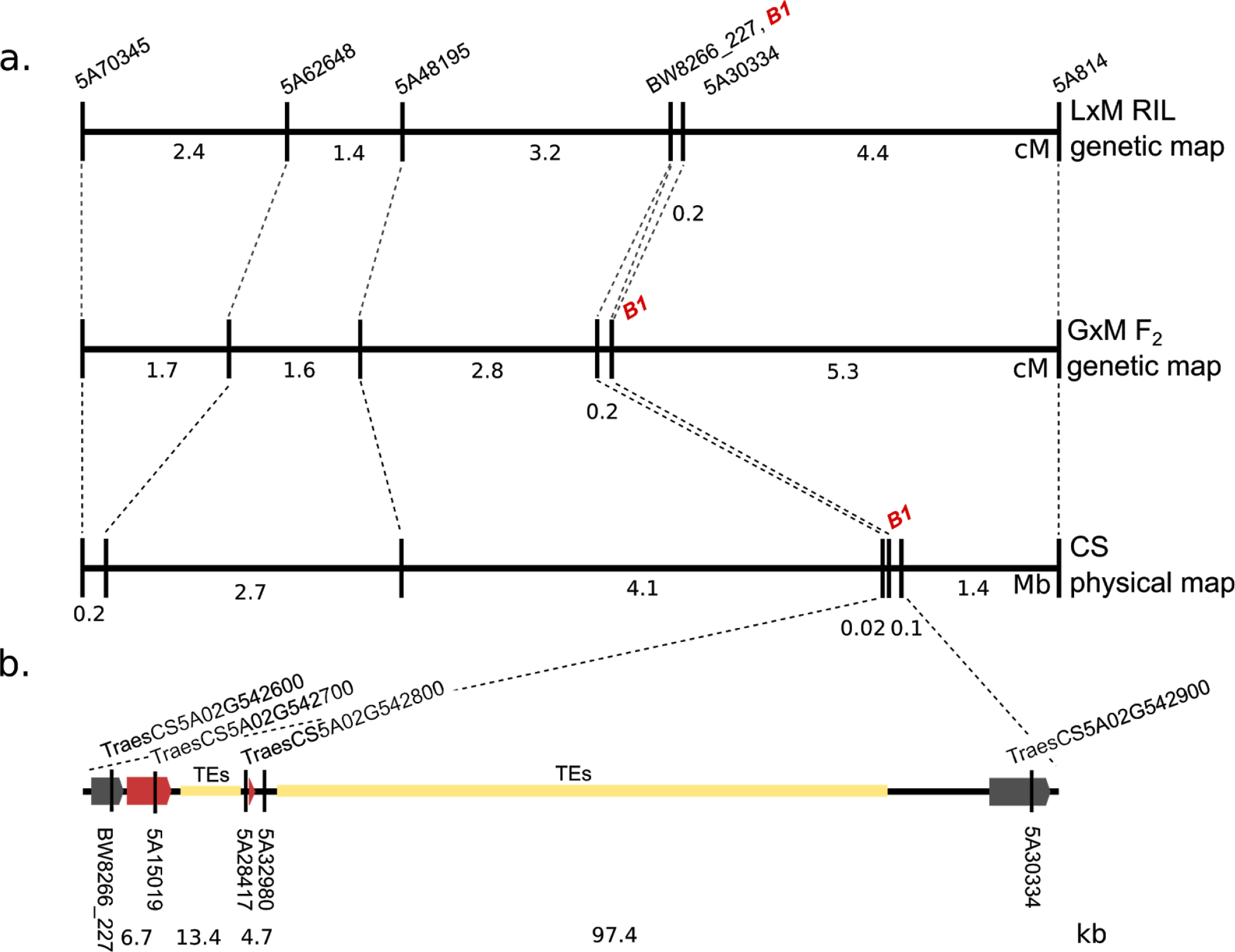
(a) Genetic distances in the *B1* region calculated from the LxM RIL population and F_2_ population GxM, compared to physical distances obtained using the IWGSC RefSeqv1.0 Chinese Spring reference genome. The *B1* locus co-segregates with SNP markers *5A15019* and *5A28417*. (b) The fine-mapped *B1* region in Chinese Spring, containing the candidate *AWNS-A1* gene (TraesCS5A02G542800), is shown relative to the genetic and physical maps. Genes co-segregating with *B1* are highlighted in red. Approximate positions of SNP markers are shown, including the most significant GBS marker in both RIL population and GWAS results (*5A28417*), and the marker most predictive of awn status in global germplasm (*5A32980*).

Analysis of awned M2 plants of the awnless variety Brundage identified deletions on the distal part of 5AL. Genomic DNA of awned and awnless M2 plants along with Chinese Spring and deletion line 5AL-6 (TA4535-6) missing the terminal 32% of 5AL was used to amplify 11 KASP markers targeting 5AL from 696 Mb to 706 Mb. For all markers, amplification was observed for wild-type Brundage, plants from each M2 family, and from Chinese Spring. No amplification was obtained for deletion line 5AL-6, suggesting genome specificity of the primers. Deletions in the region were observed for all 53 awned plants selected from 17 segregating M2 families (Table 1). The majority of the M2 lines had lost more than 7.5 Mb of the surrounding genome, with the smallest deletion being between 19.9 kb and 392 kb in size.

**Table 1.**
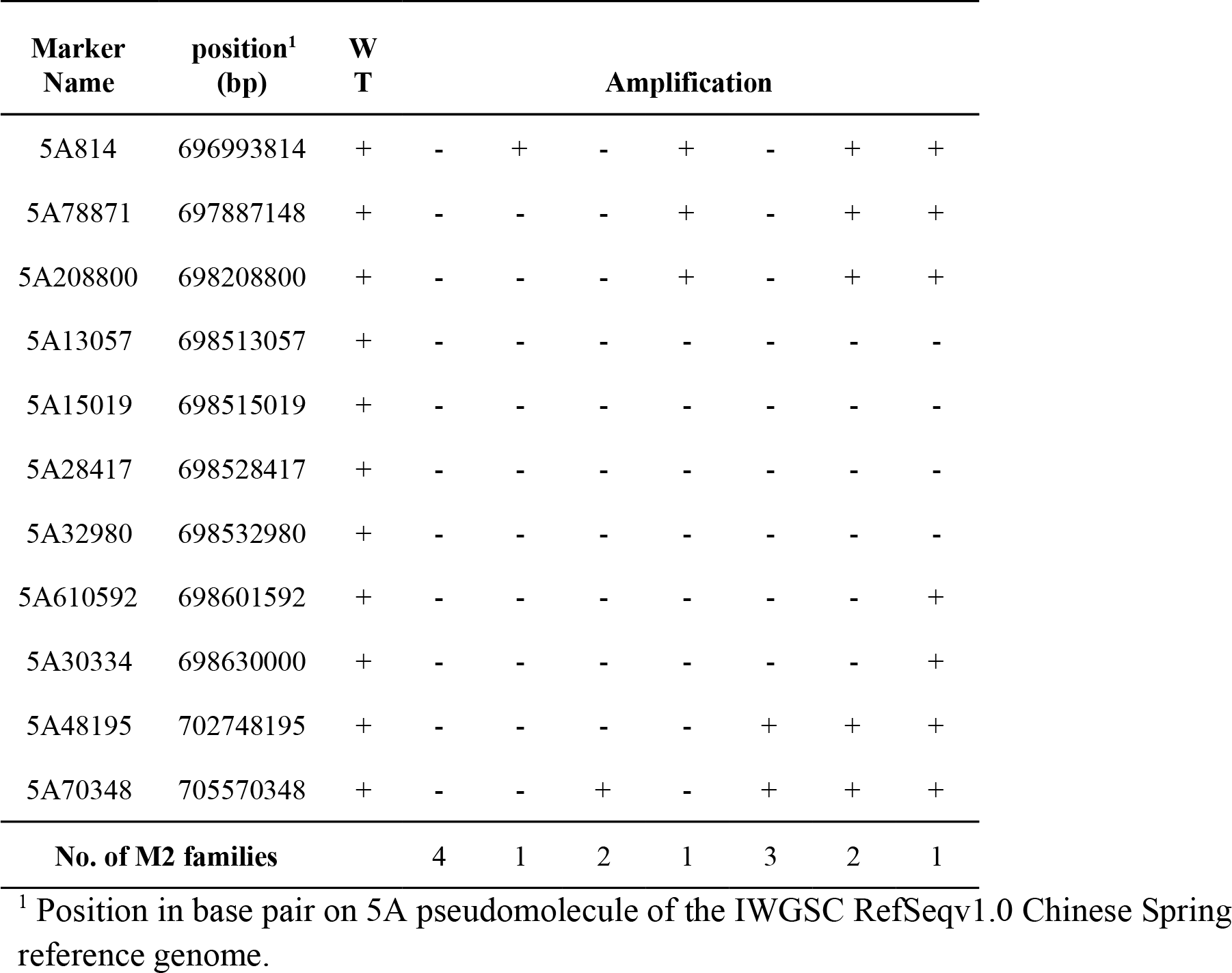
Presence (+) and absence (−) of amplification with markers flanking the *B1* locus in awned M2 families developed by fast neutron irradiation of the awnless cultivar Brundage. Number of families (unique mutation events) for each haplotype is recorded. All reported haplotypes were validated in multiple individuals from each family.

Evaluation of exome capture data identified additional SNPs between the parental lines LA95135 and SS-MPV57 upstream and within predicted gene TraesCS5A02G542700. The LA915235 and SS-MPV-57 haplotypes were compared to sequence data surrounding the orthologous gene TRIUR3_34498 from *Triticum urartu*, the wild progenitor of the A-genome in wheat. Given that accessions of *T. urartu* possess awns, a potential sequence variant underlying awn suppression would be shared by LA95135 and *T. urartu* and differ from awnless line SS-MPV57. A KASP marker was designed from a SNP unique to SS-MPV57 predicted to create a missense mutation in exon 9 (*5A16541*). The winter diversity panel was screened with this marker and the marker for an intron SNP (*5A15019*). Of the 455 lines screened, 55 individuals with the awnless *5A15019* allele and 99 individuals with the awnless *5A16541* allele were awned, suggesting that both polymorphisms are linked to the causative polymorphism underlying awn suppression but that neither are causitive themselves.

### Characterization and expression of candidate gene *AWNS-A1*

Fine mapping placed the *B1* locus in a 127 kb region of mostly repetitive sequences in the Chinese Spring reference genome containing two predicted genes, TraesCS5A02G5426700 and TraesCS5A02G5426800. Deletion of this region in M2 mutants of awnless cultivar Brundage results in the presence of awns. Marker analysis of wheat germplasm indicated that polymorphisms in predicted gene TraesCS5A02G5426700 annotated as universal stress protein family with protein kinase domain were not predictive of awn suppression. Thus, candidate gene TraesCS5A02G5426800, a 366 bp single exon predicted C2H2 zinc finger transcription factor seems a likely candidate for the *B1* awn supressor and we have named this gene *AWNS-A1*. A predicted protein sequence was obtained via Expasy translate, and blastp was used to identify homologous genes in the Uniprot database in *Arabidopsis thaliana*. The candidate gene shares conserved zinc finger and terminal ethylene-associated response motifs (EAR-like) with a family of C2H2 zinc finger developmental transcription factors in *Arabidopsis* (Fig. 4), including transcription factors associated with floral development. Variation within the coding sequence was not was found among the Chinese Spring refseq v1.0, scaffold assemblies of awnless winter wheats with the *B1* suppressor (Cadenza, Paragon, Robigus, and Claire) and the awned tetraploid wheat Kronos (https://opendata.earlham.ac.uk/opendata/data/). In Chinese Spring, *AWNS-A1* is surrounded by more than 100kb of repetitive elements, hindering assembly and comparison of the region in the scaffold assemblies. In the available awnless scaffolds, however, at least 10kb of additional repetitive elements were observed upstream of *AWNS-A1*, suggesting insertion or removal of transposable elements since the divergence of the available awnless assemblies and Chinese Spring. Despite a lack of coding sequence polymorphisms, up-regulation of a transcription factor in awnless varieties due to TE insertions or polymorphisms in regulatory regions might explain the dominance of the *B1* allele.

**Fig 4.**
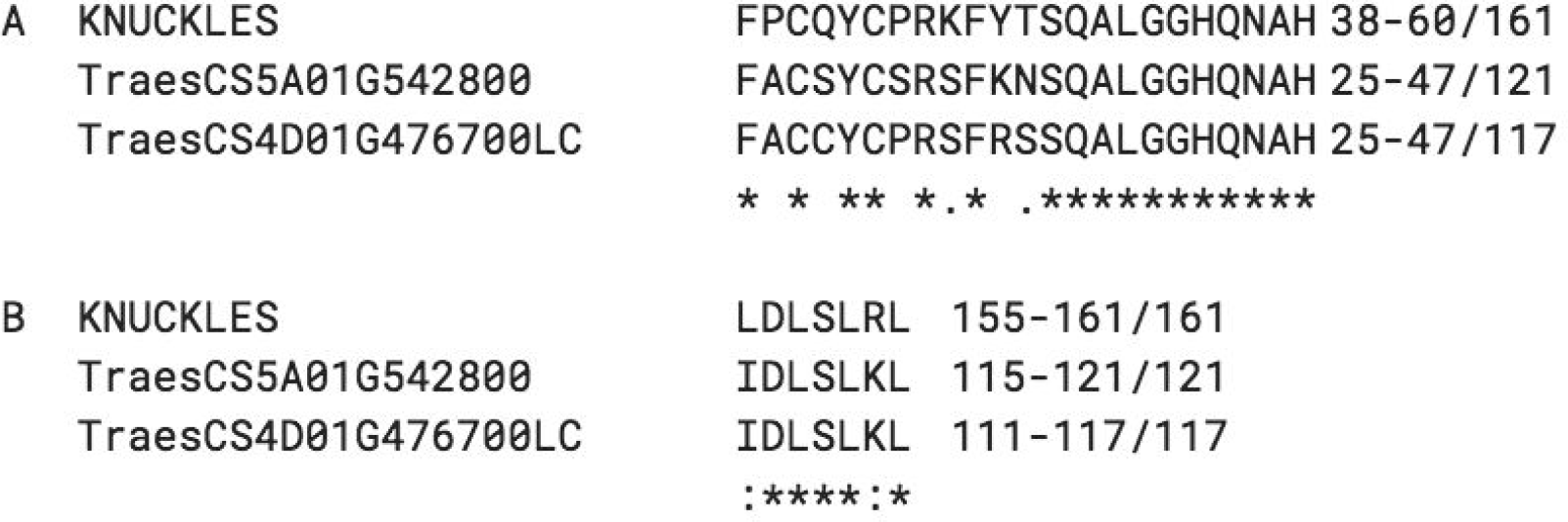
Alignment of functional motifs from characterized *Arabidopsis* C2H2 zinc finger KNUCKLES with candidate *AWNS-A1* gene (TraesCS5A01G542800) and its homoeologue *AWNS-D1* (TraesCS4D01G476700LC). KNUCKLES protein sequence downloaded from Uniprot database, candidate gene and ortholog sequences predicted via Expasy. A) Cystedeine and histidine residues responsible for zinc finger binding and core plant zinc finger sequence QALGGH are conserved between AWNS1 and KNUCKLES (Takatsuji, 1999 from Payne, Johnson, and Koltunow 2004.) B) Conserved EAR-like motifs of KNUCKLES (Hiratsu et al., 2002 from Payne *et al.*, 2004) and AWNS1.

*AWNS-A1* is located in a region of 5AL that originates from a translocation from the 4A chromosome and homoeologous AWNS1 genes were identified on the long arms of chromosomes 4B and 4D. *AWNS-B1* on 4BL contains a frameshift mutation and is not annotated in either the high confidence or low confidence gene set of the IWGSC v1.1 annotation (https://wheat-urgi.versailles.inra.fr/Seq-Repository/Annotations). The *AWNS-D1* ortholog is similar to *AWNS-A1* (Fig. 4), with *AWNS-A1* differing somewhat in the region between the zinc finger and EAR-like motifs, including a 10 amino acid insertion from positions 90-99. The WheatEXP database was used to evaluate patterns of gene expression of the AWNS1 homoelogs. *AWNS-A1* expression increases as the early inflorescence develops, and decreases after the emergence of the head (Fig. S1), while *AWNS-D1* increases in expression later in the development of the spike. *AWNS-B1* was not found in the wheat expression database. The WheatFP browser (http://bar.utoronto.ca/efp_wheat/cgi-bin/efpWeb.cgi)(Ramirez-Gonazales *et al.*, 2018, Winter *et al.*, 2007 was used to localize gene expression in an awned spring variety *Azhurnaya*. *AWNS-A1* is most highly expressed in the developing inflorescence, with some expression in the developing awns, ovaries, and grain tissues, while *AWNS-D1* is mostly expressed later in spike development (Fig. S2, S3). Similar analyses of predicted gene TraesCS5A02G5426700 having a protein kinase domain indicate that this gene is expressed in most tissues, with higher expression in spikes after the development of awn tissue (Fig. S4).

RNA was isolated from developing spikes of plants of awned cultivars LA95135 and AGS 2000 and awnless cultivars SS-MPV57 and Massey. Gene expression of *AWNS-A1* quantified through qPCR against actin showed a significant 7-fold average increase in gene expression for awnless individuals (P < 1e-7) (Fig. **5a**). As RNA sequencing data show that the gene is differentially expressed at different developmental stages, the developing spikes were grouped into four stages based on development of the meristems (Fig. **5b**). Within each group, expression of the candidate gene was higher in awnless individuals and increased at time points associated with awn development, suggesting a mechanism of awn suppression associated with *AWNS-A1*. Although polymorphisms in *AWNS-A1* were not detected in any of the germplasm analyzed, differences in expression may result from polymorphisms in cis-regulatory regions, either directly upstream of a candidate gene in the promoter or in more distant enhancers or repressors.

**Fig. 5a.**
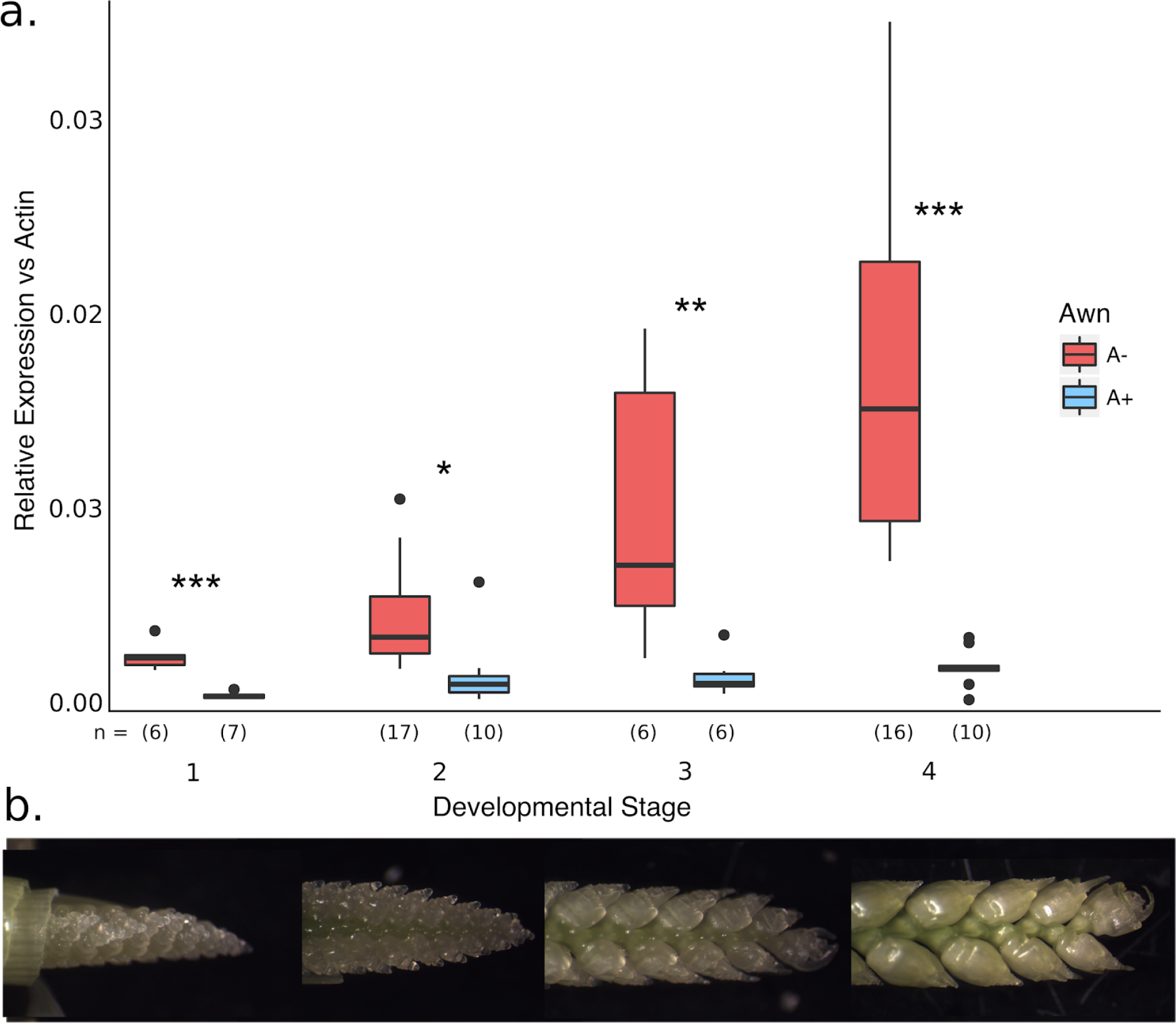
Expression of *AWNS-A1* in apical meristems of awned (blue) and awnless (red) wheat plants at different developmental stages where 1 = younger meristems and 4 = older meristems. Expression is relative to the reference gene *β-ACTIN*. Significant difference between awned and awnless individuals are indicated (*** P < .001, ** P < .01, * P <.05). Number of biological replicates (n) is given per group, with three technical replications per biological replicate. 5b. Representative spikes at each developmental stage.

### Haplotype diversity in global wheat germplasm

A 7 kb region of non-repetitive sequences surrounding the *AWNS-A1* candidate gene was delineated after BLAST against the Triticeae repeat sequences database of wheat transposable elements (https://wheat.pw.usda.gov/ITMI/Repeats/blastrepeats3.html). This region was Sanger sequenced in a set of 10 individuals, including LA95135, SS-MVP57, and eight accessions of *T. aestivum* from the NSGC wheat core collection having different haplotypes for KASP assays in the region from 698.51 Mb to 698.62 Mb of 5AL (Table S4). Contigs from each individual were aligned along with available sequences of the region from Chinese Spring, awnless wheats Cadenza, Paragon, Robigus, and Claire, the awned tetraploid wheat Kronos, and the diploid A-genome progenitor *T. urartu*. No polymorphisms in the coding sequence were observed, and sequence variation upstream of the gene in the promoter region (including marker *5A28417*) was not predictive of awn suppression in this set of cultivars. However, a 30 bp deletion 4 kb downstream of the gene was predictive of awn suppression in the alignment set.

KASP marker *5A32980* designed around this 30 bp deletion were evaluated on DNA isolated from 2439 winter and spring wheat accessions in the USDA NSGC core set. Of the 455 winter wheat accessions evaluated, 57% were awnless compared with 45% of 1984 spring wheat accessions (Table 2). The frequency of awn suppression was highest in accessions from Western Europe with 93% of winter wheat and 61% of spring accessions being awnless. KASP marker *5A32980* was the most predictive marker of awn status. Ninety-eight percent of the 196 winter wheat accessions homozygous for the non-deleted allele of marker *5A32980* had awns. The *5A32980* marker was predictive of awn suppression in all but 59 of 696 awnless spring lines (92%). The *Hd* and *B2* alleles can also suppress awns in wheat. It is possible that awnless accessions without the 30 bp deletion downstream of *AWNS-A1* may possess *Hd* or *B2* rather than *B1*. Of 263 winter wheat accessions homozygous for the deletion, all but three were categorized as awnless (98%). Of the 1295 awned spring wheats screened, 15 lines were found to possess the non-deleted *5A32980* allele. Of these, 13 accessions are landraces from Sudan, Egypt, and Oman, suggesting they may share a rare genetic variant in either the candidate gene region or at the AWNS-A1 target.

**Table 2.** Observed haplotypes in diverse wheat germplasm near *B1*. For each KASP marker, allele A is the LA95135 (*b1*) allele, allele B the MPV57 (*B1*) allele, and allele C a null allele. Position of SNPs on 5A pseudomolecule of IWGSC Chinese Spring RefSeqv1.0 are indicated below the marker name.

| Haplotype | Marker Name (position<sup>1</sup>) |  |  |  |  |  |  | Assoc.<br>allele |
| --- | --- | --- | --- | --- | --- | --- | --- | --- |
|  | 5A15019 | 5A28417 | 5A29396 | 5A32980 | 5A91913 | 5A613482 | 5A614919 |  |
|  | (698515019) | (698528417) | (698529396) | (698532980) | (698591913) | (698613482) | (698614919) |  |
| Hap1 | A | A | A | A | A | A | A | b1 |
| Hap2 | A | B | A | A | A | A | A | b1 |
| Hap3 | B | B | A | A | B | A | B | b1 |
| Hap4 | B | B | A | A | B | B | A | b1 |
| Hap5 | B | B | A | A | B | B | B | b1 |
| Hap6 | B | B | C | A | B | B | C | b1 |
| Hap7 | B | A | B | B | B | B | B | B1 |
| Hap8 | B | B | B | B | B | B | B | B1 |

<sup>1</sup> Position in base pair on 5A pseudomolecule of the IWGSC RefSeqv1.0 Chinese Spring reference genome.

Eight haplotypes were identified when six SNP markers flanking *AWNS-A1* were examined in conjunction with marker *5A32980*. Marker haplotype 1 (Hap1) present in the awned cultivar LA95135 and Hap8 present in awnless cultivar SS-MPV57 were the most common, detected in 55% and 35% of accessions, respectively. Greater diversity in haplotypes was observed in spring accessions as only four of the eight haplotypes were detected in winter wheat accessions. The greatest diversity in haplotypes was in accessions from Central Asia where all haplotypes expect Hap4 were observed. Although all eight haplotypes were present in accessions from Western Europe, 95% of these accessions had either Hap1 or Hap8. Presence of haplotypes were related geographic origin. For example, Hap6 was present primarily in accessions from Eastern Europe (16%) and Hap3 was most common in accessions from Central Asia (5%).

These results suggest that the *B1* locus is present in the vast majority of awnless winter wheat (98%) and most awnless spring wheats (92%). Awnless individuals carrying the *b1* allele likely contain some combination of the *Hd* and *B2* alleles as in Chinese Spring. Of these lines, 67% originated from central and south Asia, primarily Nepal and India, suggesting regional variation in control of awn suppression. However, regional biases in the initial selection of the diversity panels (for example, only four awnless spring wheat lines from China were included) limit the conclusions that can be drawn about geographic distribution of the three major awn suppression alleles.

## Discussion

Wheat’s large genome size, hexaploidy, and long range linkage disequilibrium can make association mapping more challenging than in other species. In this study, association mapping in a set of elite lines identified a highly significant SNP maker for awn suppression. Alignment of reads from the reduced representation libraries developed for GBS to the recently annotated and assembled reference sequence identified a significant SNP nearby a candidate gene encoding a C2H2 zinc finger transcription factor. Targeted mapping in large bi-parental mapping populations determined the significant marker co-segregated with awn suppression. Further analysis using fast neutron mutants, a panel of wheat lines representing global wheat diversity, and gene expression in developing spikes support the identification of this C2H2 zinc finger transcription factor, designated *AWNS-A1*, as the causal gene underlying the *B1* awn suppressor. To our knowledge, this is the first case of association mapping using a GBS marker data set resulting in the isolation of an agronomically important gene in wheat.

Global variation for awn length suggests that the presence or absence of awns may be differentially adaptive to varying environments and production systems. Awns are known to contribute to yield in warmer, drier environments, cooling the wheat spike and supplying carbohydrates to developing grain. (Grundbacher 1963, Kjack & Witters, 1974, Motzo & Giunta, 2002, Li *et al.* 2006, Tambussi *et al.* 2007, Ali *et al.* 2010, Maydup *et al.* 2014). Identification of a significant association of the *B1* awn suppressor with reduced test weight in our association panel suggests that awns influence kernel size in winter wheat grown in the southeastern United States. In the biparental LM RIL population, QTL analysis confirmed that the presence of awns contributed to increase kernel weight. In the current study, evaluation of kernel morphometric traits also determined that the increased kernel weight in lines with the *b1* allele was associated with greater kernel length.

Except in forage cultivars, the most economically important organ in wheat is the spike. As such, developing a better understanding of the gene networks that control spike development is critical for wheat breeders. Consistent with other studies, we observed that awn suppression was associated with an increase in the number of spikelets per spike (Rebetzke *et al.*, 2016). A major hindrance to increasing yield of modern wheat cultivars is the availability of sink tissues to fill with carbohydrates -- increasing spikelet number is therefore one strategy to increase the number of grains harvested per unit area (Miralles & Slafer 2007). The final number of rachis nodes in wheat is correlated with the duration of the reproductive growth period, and genes associated with flowering time in wheat may influence the number of spikelets per spike. In our QTL mapping study the segregating *Ppd-D1* and *Rht-D1* genes were associated with QTL for both spikelets per spike and flowering time, but *B1* was only associated with spikelets per spike, indicating that it influences the number of rachis nodes through a different mechanism. The trade-off between increased number of grains per spike and reduced kernel size observed in this study is well documented (Sadras 2007). New research into control of grain size in wheat has identified genes such as the grain weight gene, TaGW2, a negative regulator of grain width. Knock down mutations in TaGW2 homoeologs additively increase grain size as each homeologous copy is disabled. Improved understanding of the control of spikelets per spike, grain number and grain size in wheat will provide insight in to pathways that can be manipulated to increase grain yield. Identification of the *AWNS-A1* transcription factor allows us to design further experiments to identify downstream targets and effects of this gene, and to begin exploring its interaction with other floral developmental genes.

While the simple inheritance and obvious phenotype created by the *B1* locus facilitate selection for awned plants, a predictive marker for awn supression should help wheat breeders introgress the *b1* allele into awnless cultivars, and vice versa. Wheat breeders increasingly rely on genomic prediction, in which whole-genome marker data is used to estimate breeding values. Awns have a significant effect on test weight in our environment, and are identified as a major QTL for number of spikelets per spike, a yield component trait, in our biparental population. As genomic prediction relies on linkage between markers and causal variants, combining GBS data with predictive markers for major-effect QTL should improve prediction accuracy. Given that the *B1* locus is the sole determinant of awn status in the southeastern U.S. and likely the major determinant of awn status globally, a predictive marker can be used to improve accuracy of genomic prediction models in wheat.

The *AWNS-A1* gene is orthologous to a set of zinc finger transcription factors in *Arabidopsis*, all of which contain a zinc finger and EAR-like motif. This family of transcription factors are usually repressors; the zinc finger domain binds to the target sequence, and the EAR-like motif recruits histones which down-regulate the target gene (Kagale & Rozwadowski 2010). The best characterized member of this family, KNUCKLES (KNU), helps terminate stem cell maintenance in *Arabidopsis* (Payne *et al.*, 2004). The floral homeotic protein AGAMOUS (AG) up-regulates KNU by binding to its promoter, with KNU in turn repressing the homeodomain protein WUSCHEL (WUS) until a specific developmental stage (Sun *et al.*, 2009). WUS is responsible for the maintenance of stem cells in *Arabidopsis*, and over-expression of KNU preemptively terminates flora meristem development (Sun *et al.*, 2009). In this way, KNUCKLES represses growth of certain floral tissues to allow other floral tissues to develop at the same pace. KNU expression in cells reduces their proliferation (Payne *et al.*, 2004). If *AWNS-A1* plays a similar role in wheat, its overexpression suppressing developing awn tissue suggests a potential explanation for the dominance of the *B1* allele.

In a biparental mapping population between Chinese Spring (*Hd b1 B2*) and Courtot (*hd b1 b2*), Sourdille *et al.* (2002) observed no QTL for awn length near the *B1* locus but noted a slight awned phenotype in a 5AL deletion line, concluding that Chinese Spring’s genotype was (*Hd B1 B2*). Sequence data indicates that Chinese Spring has the *b1* allele; however, we observed a similar phenotype in the deletion line 5AL-6 (TA4535-6) (Fig. S6). The small awns in the deletion line phenotype might instead be explained by a lower dosage of *AWNS-A1* compared to wildtype Chinese Spring (Fig. S5). The interaction between the three awn suppressors suggests that they are involved in the same pathway. While all three awn suppression genes are treated as dominant genes, intermediate phenotypes are sometimes observed in plants heterozygous for the *B1* allele. Past studies have treated these intermediate forms as having a genetic origin in either an additional “half-awn” gene (A), or as an additional allele at the *B1* locus (*B1*^a^) (Watkins and Ellerton, 1940). We observe variation for awn length within awned individuals in both spring and winter diversity panels, including lines approaching a “half-awn” phenotype. However, we also observe variation for awn length in F4 RIL families and F2 populations segregating for the *B1* allele. It seems likely that “half-awn” phenotypes can be produced by either additional genes controlling awn length or by heterozygosity at the *B1* locus.

Transposable elements, particularly long terminal-repeat retrotransposons (LTR-RTs), are the dominant feature of plant genomes (Parisod *et al.*, 2010). This is especially true of many grass species, with TEs composing roughly 85% of the maize and wheat genomes (Appels *et al.*, 2018, Schnables *et al.*, 2009). Our understanding of the mechanisms by which transposable elements shape plant evolution continues to grow. The expansion and contraction of the plant genome mediated by TEs influences the character of adaptive variation, individual TE insertions may disrupt or alter methylation patterns of nearby genes, and TEs may be co-opted to provide new functions (Mei *et al.*, 2018, Wittmeyer *et al.*, 2018, Studer *et al*., 2011, Lisch 2009). Alignment of *AWNS-A1* gene to homeologous regions in the B and D genomes reveal major variation in transposable elements both upstream and downstream of the candidate gene. In addition, preliminary assemblies of the region in wheat lines carrying the *B1* allele reveal further TE expansion in the region compared to lines carrying the *b1* allele. Given the importance of TE activity in functional variation, this expansion could explain the difference in expression we observe in our candidate gene between awned and awnless varieties.

Our haplotype analysis of global wheat germplasm identified two dominant haplotypes in the region of non-repetitive sequence flanking *AWNS-A1*, with a smaller number of lines having haplotypes associated with geographic regions (Tables 2, 3). A 30 bp deletion 3 kb downstream of the candidate gene is predictive of the *B1* awn suppressor in nearly all lines. The region containing the 30 bp deletion is predicted to be a CpG island, where relatively higher CG content suggests a lack of methylation-mediated C to T conversions. The deleted sequence also contains motifs associated with transcription factor binding, but the ability of software to predict transcription factor binding from sequence data is limited. Determining if the observed up-regulation of the candidate gene is a product of said deletion, a byproduct of TE insertions, or some other causative polymorphism will therefore require further work. However, given the observed linkage disequilibrium between the predictive deletion and phenotype, the difference is inconsequential for practical applications of the marker.

**Table 3.** Number of lines having observed haplotypes in spring and winter wheat germplasm based on geographic region of origin for accessions. For each haplotype, left column represents number of awnless lines (A-) and the right column the number of accessions having awns (A+).

| Region of origin | Growth habit | Hap1 |  | Hap2 |  | Hap3 |  | Hap4 |  | Hap5 |  | Hap6 |  | Hap7 |  | Hap8 |  | Total |
| --- | --- | --- | --- | --- | --- | --- | --- | --- | --- | --- | --- | --- | --- | --- | --- | --- | --- | --- |
|  |  | A- | A+ | A- | A+ | A- | A+ | A- | A+ | A- | A+ | A- | A+ | A- | A+ | A- | A+ |  |
| East Asia | Spring | 2 | 43 | - | - | - | 2 | - | 1 | - | - | - | - | - | - | 13 | - | 61 |
|  | Winter | - | 15 | - | - | - | - | - | - | - | - | - | - | - | - | 4 | - | 19 |
| South Asia | Spring | 39 | 58 | 2 | 2 | - | 3 | - | - | - | - | - | - | - | - | 12 | - | 116 |
| Caucuses | Spring | 4 | 52 | - | - | - | 1 | - | - | - | 2 | - | - | - | - | 20 | - | 79 |
|  | Winter | 1 | 11 | - | - | - | - | - | - | - | - | - | - | - | - | 4 | - | 16 |
| Central Asia | Spring | 15 | 193 | - | 2 | - | 13 | - | - | - | 9 | - | 1 | 1 | - | 70 | - | 304 |
|  | Winter | 1 | 45 | - | - | - | 8 | - | - | - | - | - | 5 | - | - | 28 | 1 | 88 |
| Western Europe | Spring | 5 | 81 | - | 1 | - | 2 | - | 1 | - | 1 | - | 3 | 3 | - | 119 | - | 216 |
|  | Winter | - | 3 | - | - | - | 1 | - | - | - | - | - | 4 | - | - | 107 | - | 115 |
| Eastern Europe | Spring | 2 | 46 | - | - | - | 3 | - | 3 | - | - | - | 10 | 1 | - | 48 | - | 113 |
|  | Winter | - | 42 | - | - | - | - | - | - | - | - | 1 | 31 | - | - | 71 | 2 | 147 |
| Middle East | Spring | 6 | 156 | - | 3 | - | 2 | - | - | - | - | - | - | 1 | - | 38 | 4 | 210 |
|  | Winter | - | - | - | - | - | - | - | - | - | - | - | 1 | - | - | 1 | - | 2 |
| Sub-Saharan Africa | Spring | 8 | 149 | 1 | 3 | - | 1 | - | 1 | - | - | - | - | 1 | - | 79 | - | 243 |
|  | Winter | - | 1 | - | - | - | - | - | - | - | - | - | - | - | - | 1 | - | 2 |
| North Africa | Spring | 5 | 104 | - | 1 | - | 1 | - | - | - | - | - | 2 | - | - | 30 | 10 | 153 |
| North America | Spring | 5 | 47 | - | - | - | - | - | - | - | - | - | 7 | - | - | 35 | - | 94 |
|  | Winter | - | 15 | - | - | - | - | - | - | - | - | - | 1 | - | - | 11 | - | 27 |
| Oceania | Spring | - | 6 | - | - | - | - | - | - | - | - | - | - | 2 | - | 38 | - | 46 |
|  | Winter | - | - | - | - | - | - | - | - | - | - | - | - | - | - | 4 | - | 4 |
| South America | Spring | 3 | 191 | - | 4 | - | 4 | - | 1 | - | - | - | - | 1 | - | 58 | 1 | 263 |
|  | Winter | - | 5 | - | - | - | 1 | - | - | - | - | - | - | - | - | 29 | - | 35 |
| Central America | Spring | 1 | 61 | - | 1 | - | 2 | - | - | - | - | - | - | 1 | - | 20 | - | 86 |
| **Total** |  | 97 | 1324 | 3 | 17 | - | 44 | - | 7 | - | 12 | 1 | 65 | 11 | - | 840 | 18 | 2439 |

Using next-generation sequencing technologies, wheat geneticists have created communal resources that facilitate fine mapping and identification of candidate genes. The newly published wheat reference genome, tools that make it accessible to other researchers, and technologies that take advantage of the reference genome (such as gene expression and exome capture data sets) were all used in this study to reduce the time and cost of positional cloning. One of the strengths of these resources is that they become more useful as new studies utilizing them collect and contribute data. As our understanding of the genetic systems underlying complex traits increases, these tools should facilitate the application of this knowledge to real-world crop improvements.

## Acknowledgements

The authors thank Jorge Dubcovsky and lab members for use of instruments for phenotyping grain traits. Support was provided by the Agriculture and Food Research Initiative Competitive Grant 67007-25939 (WheatCAP-IWYP) from the USDA NIFA.

