## Supplemental Information for "Sequence based mapping identifies *AWNS1*, a candidate transcription repressor underlying awn suppression at the *B1* locus in wheat"

### Supporting Information

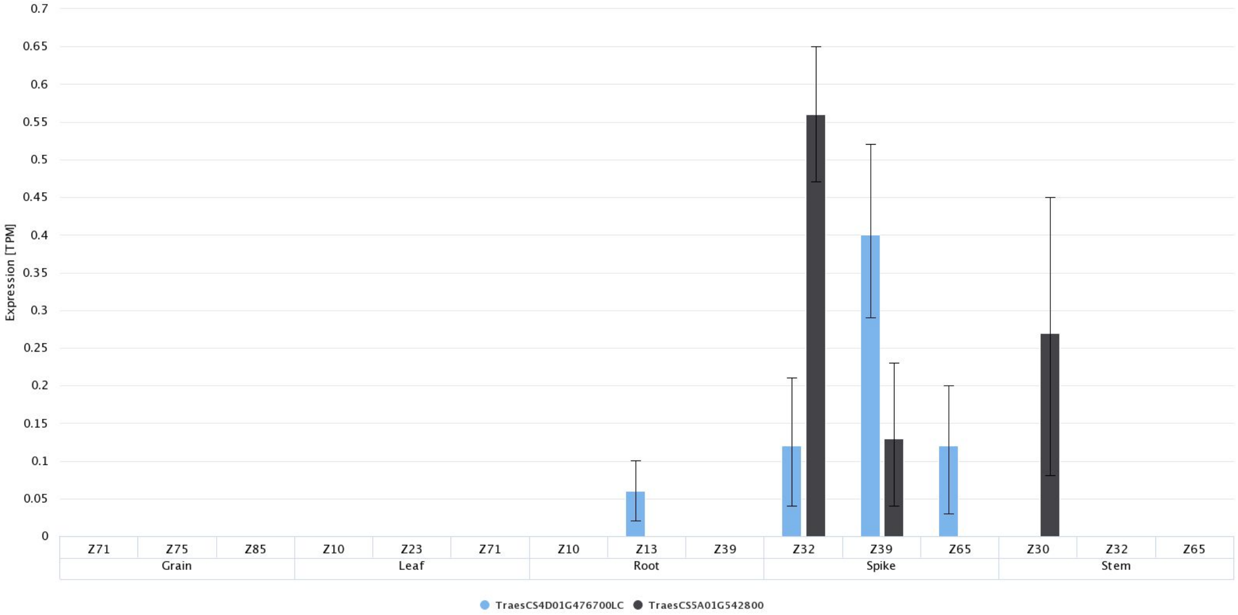


Supporting information fig. S1. Expression of *AWNS-A1* (TraesCS5A02G542800) and *AWNS-D1* (TraesCS4D01G476700LC) at three developmental time points in different tissues.

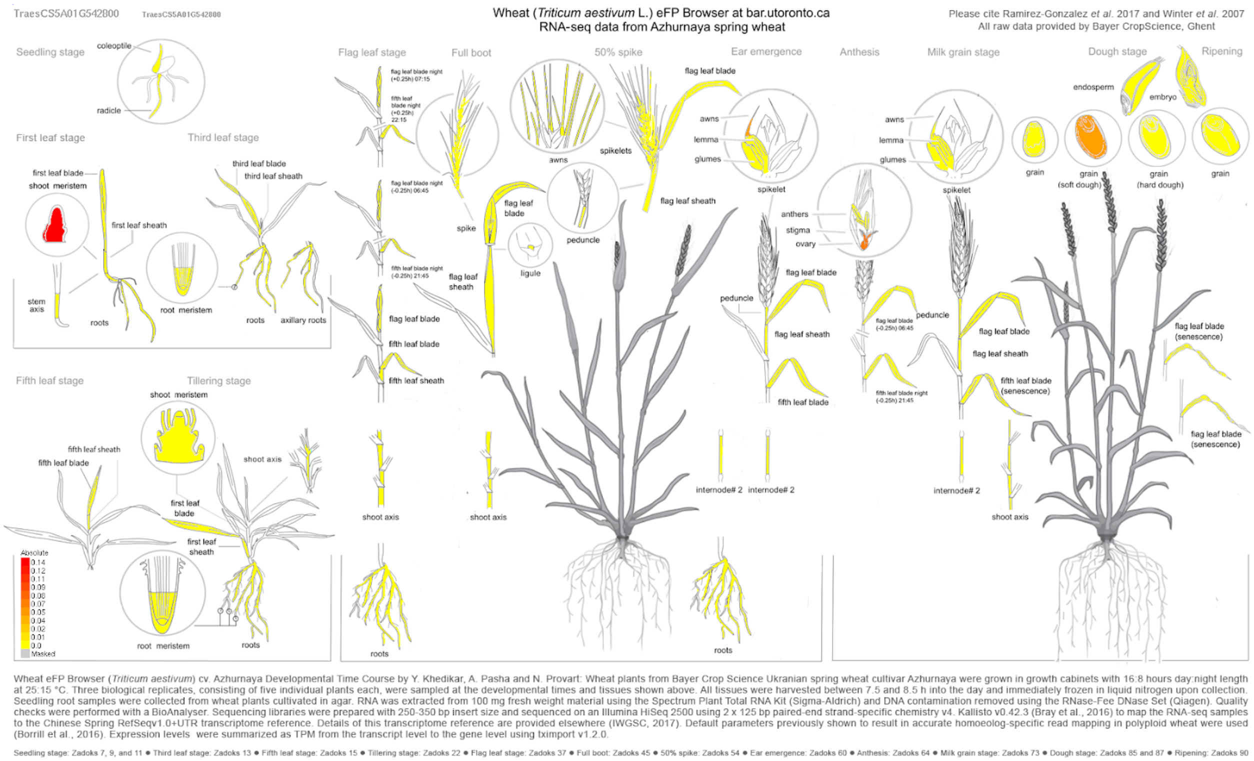


Supporting information fig. S2. Wheat eFP browser output displaying regions with no or low expression of *AWNS-A1* (yellow), and regions with higher expression (orange to red) in the awned spring wheat *Azhurnaya*. *AWNS-A1* is primarily expressed within the developing spike, as well as expressed in the awns, developing grain, and ovary.

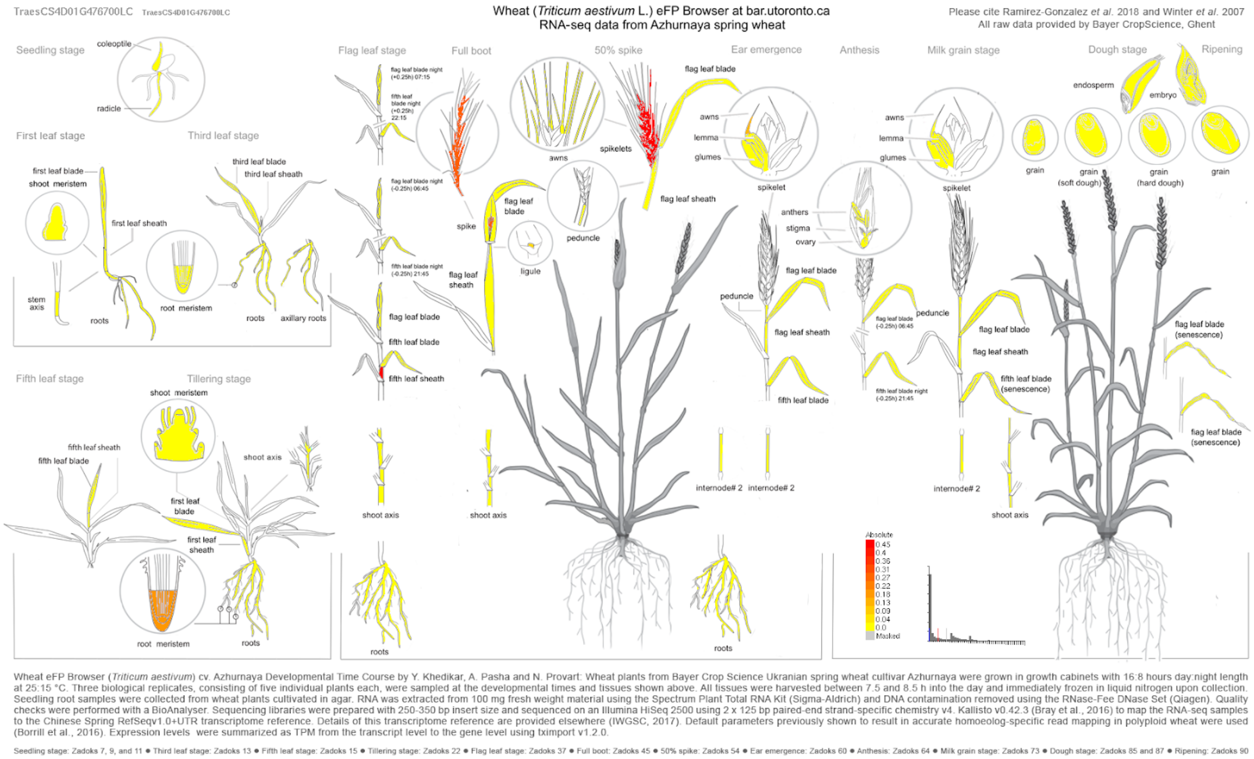


Supporting information fig. S3. Wheat eFP browser output displaying regions with no or low expression of *AWNS-D1* (yellow), and regions with higher expression (orange to red) in the awned spring wheat *Azhurnaya*. *AWNS-D1* is primarily expressed in the spike after the development of awn tissue.

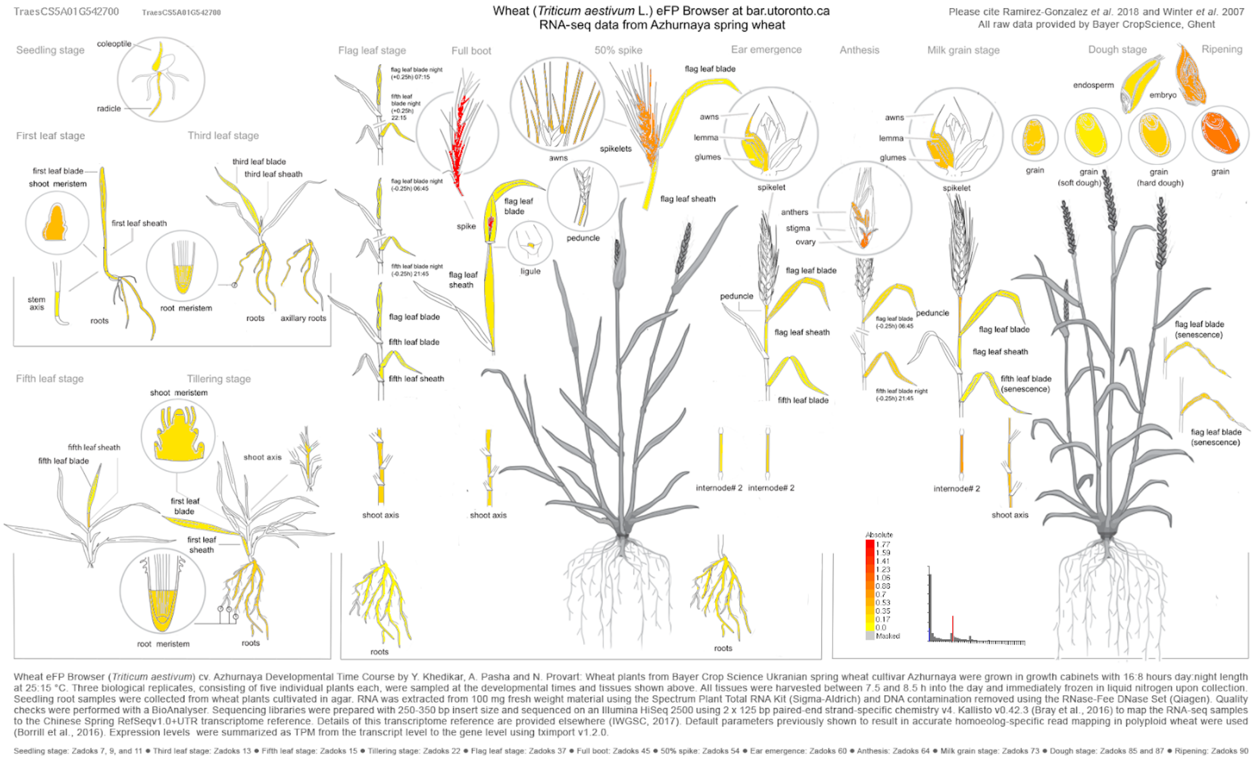


Supporting information fig. S4. Wheat eFP browser output displaying regions with no or low expression of neighboring gene TraesCS5A01G542700 (yellow), and regions with higher expression (orange to red) in the awned spring wheat *Azhurnaya*. TraesCS5A01G542700 is expressed in most tissues, with higher expression in the spike and in grain tissues after the development of awns.

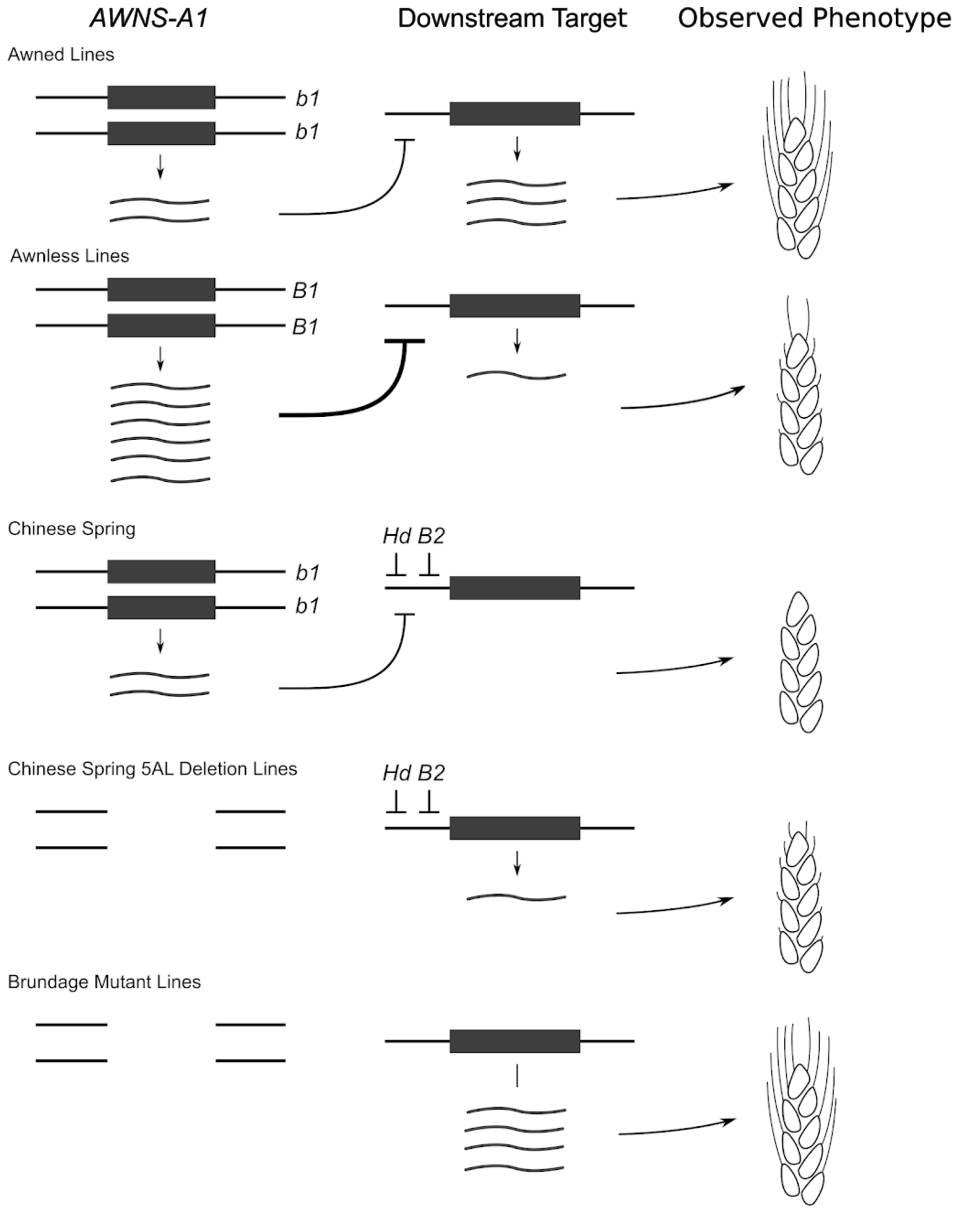


Supporting information fig. S5. Proposed mechanism of *B1* awn suppression illustrated by different allelic combinations. In wild-type *hd b2* lines, over expression of *AWNS-A1* might repress expression of a target gene controlling awn length. In Chinese Spring, *Hd* and *B2* alleles in combination with the *b1* allele result in an awnless phenotype. A 5AL Chinese Spring deletion line with the same alleles but missing *b1* displays short awn growth, while mutant Brundage lines with an *hd b2* background missing *b1* are fully awned.

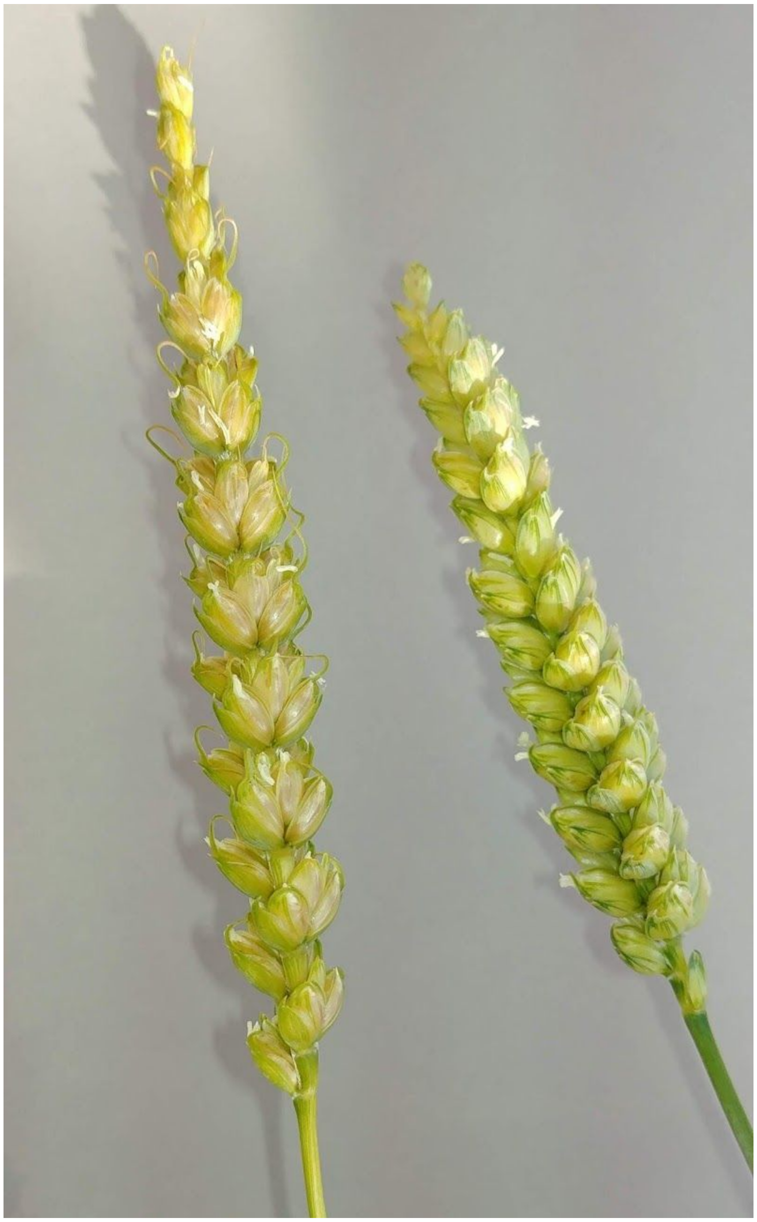

Supporting information fig. S6. Wild-type Chinese Spring (right) compared to a 5AL Chinese Spring deletion line. The elongated spike is due to the additional deletion of the nearby *Q* gene. Deletion of the wild-type *b1* allele (*Hd b1 B2*) produces small, curved spikes despite the presence of the *Hd* and *B2* awn suppressors, suggesting that all three genes may be part of the same developmental pathway.

Supplemental information table S1. Summary of markers used in this study.

| Name | SNP Position | Type | Source | SNP | Gene Id | Annotated Functional Domains |
|---|---|---|---|---|---|---|
| 5A814 | 696993814 | Intergenic | GBS | G/C | NA | NA |
| 5A78871 | 697887148 | Intron variant | Exome capture | G/A | TraesCS5A01G540400 | Formyl transferase |
| 5A208800 | 698208800 | Intron variant | Exome capture | A/C | TraesCS5A01G541400 | SNF2 family domain, Helicase conserved domain |
| BW8226_227 | 698508163 | Missense exon 2 | Illuminia SNP array (Mackay et. al 2014) | T/G | TraesCS5A01G542600 | Sugar -and other - Transporter |
| 5A13057 | 698513057 | Intron variant | Exome Capture | T/G | TraesCS5A01G542700 | Universal stress protein family, protein kinase |
| 5A15019 | 698515019 | Intron variant | Exome Capture | A/G | TraesCS5A01G542700 | Universal stress protein family, protein kinase |
| 5A16541 | 698516541 | Missense exon 9 | Sequence Alignment | A/G | TraesCS5A01G542700 | Universal stress protein family, protein kinase |
| 5A28417 | 698528417 | Upstream | GBS | T/C | TraesCS5A01G542800 | C2H2 Zinc Finger |
| 5A29396 | 698529396 | Downstream | Sequence alignment | G/A | TraesCS5A01G542800 | C2H2 Zinc Finger |
| 5A32980 | 698532980 | Intergenic | Sequence alignment | Del | NA | NA |
| 5A91913 | 698591913 | Intergenic | Sequence alignment | C/T | NA | NA |
| 5A610592 | 698610592 | Intergenic | Sequence alignment | A/G | NA | NA |
| 5A613482 | 698613482 | Intergenic | Sequence alignment | AA/CT | NA | NA |

| | | | | | | |
|---|---|---|---|---|---|---|
| 5A614919 | 698614919 | Intergenic | Sequence alignment | Del | NA | NA |
| 5A30334 | 698630334 | Synonymous variant | Exome capture | C/T | TraesCS5A01G542900 | F-box-like, Armadillo -beta-catenin-like repeat |
| 5A48195 | 702748195 | Upstream | Exome capture | T/G | TraesCS5A01G548800 | Helicase domain, P-loop_NTPase superfamily, EBV_NA3 super family |
| 5A70348 | 705570348 | Missense | Exome capture | C/G | TraesCS5A01G552800 | Domain of unknown function - DUF 4220 -, DUF594 |

Supporting information table S2. Sequences for all three primers for each KASP marker used in this study.

| Name | Allele | Sequence |
|---|---|---|
| 5A814 | A1 | GAAGGTGACCAAGTTCATGCTAAATTCCTGCTAGACACTCGTGAG |
| | A2 | GAAGGTCGGAGTCAACGGATTAAATTCCTGCTAGACACTCGTGAC |
| | C | GTTTGCGCGAATCCTGCAGCGAA |
| 5A78871 | A1 | GAAGGTGACCAAGTTCATGCTGTGCCGCCTCCATTTCCCAC |
| | A2 | GAAGGTCGGAGTCAACGGATTCGTGCCGCCTCCATTTCCCAT |
| | C | CCGTCGAACAGGTGTCGGAGC |
| 5A208800 | A1 | GAAGGTGACCAAGTTCATGCTCCTTCCCCGCTGCCCGA |
| | A2 | GAAGGTCGGAGTCAACGGATTCCTTCCCCGCTGCCCGC |
| | C | GTGCTCAAGGAGAAGCTAAGAGGTA |
| 5A13057 | A1 | GAAGGTGACCAAGTTCATGCTGGTCGATCCATCCATTCATTCATGA |
| | A2 | GAAGGTCGGAGTCAACGGATTGTCGATCCATCCATTCATTCATGC |
| | C | CCGTTTCATGTTCTAGACTGAATCTGAAT |
| 5A15019 | A1 | GAAGGTGACCAAGTTCATGCTATGAACATCTTCATTGAAACTATATTTACACA |
| | A2 | GAAGGTCGGAGTCAACGGATTGAACATCTTCATTGAAACTATATTTACACG |
| | C | GAACCCTGCAATTAACCAAAACAAAAGCTA |
| 5A28417 | A1 | GAAGGTGACCAAGTTCATGCTTGCGGCCGCTCAG |
| | A2 | GAAGGTCGGAGTCAACGGATTGCATGCTTGCGGCCGCTCAA |
| | C | GACAGTAATAATGCTGCAGTAGATGTGTA |
| 5A29396 | A1 | GAAGGTGACCAAGTTCATGCTGCCTAGAGACAAAAATAAAGTTATTATTATTC |
| | A2 | GAAGGTCGGAGTCAACGGATTGGCCTAGAGACAAAAATAAAGTTATTATTATTT |
| | C | ACCATTGCAATTATAGCACCAAGATATAAA |
| 5A32980 | A1 | GAAGGTGACCAAGTTCATGCTAGCTACGGGCCCACTTRGACA |
| | A2 | GAAGGTCGGAGTCAACGGATTCTACGGGCCCACTTRGACG |

| | | |
|---|---|---|
| | C | CCTGCGGGGCTCCCAGCAA |
| 5A91913 | A1 | GAAGGTGACCAAGTTCATGCTCACATGCAACACACCACTTGTCA |
| | A2 | GAAGGTCGGAGTCAACGGATTCACATGCAACACACCACTTGTCG |
| | C | TTCACACTCCTACTTCCCCAGGTT |
| 5A613482 | A1 | GAAGGTGACCAAGTTCATGCTTTCCAGCAAAGTTGGAAGTACAATTTA |
| | A2 | GAAGGTCGGAGTCAACGGATTCCAGCAAAGTTGGAAGTACAATTTC |
| | C | TAGTAAAGCGCGTCCAACCCTTCTA |
| 5A614989 | A1 | GAAGGTGACCAAGTTCATGCTGATTAAGATATTCAATTTTGGATTTGATTCATA |
| | A2 | GAAGGTCGGAGTCAACGGATTAAGATATTCAATTTTGGATTTGATTCATG |
| | C | GATCAAAATTGAAAGCGTTGATCTGGTCAA |
| 5A610592 | A1 | GAAGGTGACCAAGTTCATGCTATACACGGCTTTCCACAATTAGTTGT |
| | A2 | GAAGGTCGGAGTCAACGGATTACACGGCTTTCCACAATTAGTTGC |
| | C | CCCTAACTAACATATAGCCATGTGCAAT |
| 5A30334 | A1 | GAAGGTGACCAAGTTCATGCTCTTCGAGTGGCAAGCGCAAC |
| | A2 | GAAGGTCGGAGTCAACGGATTCTCTTCGAGTGGCAAGCGCAAT |
| | C | CCATGTGGGTTGGTCTCAGGGAT |
| 5A48195 | A1 | GAAGGTGACCAAGTTCATGCTGGTTTCTTCAGAAAATGGAGGTCGA |
| | A2 | GAAGGTCGGAGTCAACGGATTGTTTCTTCAGAAAATGGAGGTCGC |
| | C | CTTCATTGATGACCTCCCCATCTTTATAT |
| 5A70348 | A1 | GAAGGTGACCAAGTTCATGCTGCGGTCTTGCAGGATGCTGAC |
| | A2 | GAAGGTCGGAGTCAACGGATTGCGGTCTTGCAGGATGCTGAG |
| | C | GGGGTAGCCATACTTCCTCTATAGAT |
| BW8226_227 | A1 | CGTCCATGGAGTCGTTCCTCAAT |
| | A2 | GTCCATGGAGTCGTTCCTCAAG |
| | C | GTGGTACACGTCCGGGAAGAAT |

Supporting information table S3. Line names, observed haplotypes and geographic origin of accessions selected for sequencing of the AWNS-A1 gene and surrounding region.

| Name | Haplotype | Origin | Awns |
|---|---|---|---|
| SS_MPV-57 | Hap8 | Virginia | - |
| LA-95135 | Hap1 | Louisiana | + |
| AR05055-1-1 | Hap3 | Arkansas | + |
| Echinoides | Hap3 | Afghanistan | + |
| Ideal | Hap3 | Denmark | + |
| Sandu 22 | Hap6 | Romania | + |
| Tadzyksaja 13 | Hap8 | Tajikstan | - |
| Winter Bearded | Hap8 | UK | - |
| IWA 8613594 | Hap8 | Iran | - |
| IWA 8606785 | Hap3 | Iran | + |

Supporting information table S4. Sequences for qPCR primers used for quantifying gene expression.

| Name | Sequence |
|---|---|
| Actin-SYBR-F1 | ATGGAAGCTGCTGGAATCCAT |
| Actin-SYBR-R1 | CCTTGCTCATACGGTCAGCAATAC |
| AWNSA1_24_F | GGAGATGGAAGAGGGGCTCGAT |
| AWNSA1_8_R | TTGAAGCTGCGTGAGCAGTAGG |

Supporting information table S5. Genome position and LOD scores of markers significantly associated with presence or absence of awns in association mapping panel of 640 soft red winter wheat lines.

| SNP | maf | |
|---|---|---|
| S5A_681455268 | 0.163964 | 0.000321 |
| S5A_684942630 | 0.320721 | 3.43E-07 |
| S5A_685141655 | 0.368468 | 7.79E-06 |
| S5A_690417237 | 0.083784 | 1.56E-09 |
| S5A_693326887 | 0.227027 | 0.000183 |
| S5A_695581531 | 0.451351 | 2.75E-08 |
| S5A_696479493 | 0.477477 | 0.000153 |
| S5A_696993814 | 0.446847 | 1.11E-08 |
| S5A_697552811 | 0.266667 | 1.56E-09 |
| S5A_697590026 | 0.252252 | 7.37E-08 |
| S5A_698003176 | 0.283784 | 3.21E-06 |
| S5A_698127281 | 0.285586 | 6.38E-07 |
| S5A_698225912 | 0.263964 | 1.05E-07 |
| S5A_698320453 | 0.257658 | 2.35E-08 |
| S5A_698528417 | 0.404505 | 7.19E-57 |
| S5A_699803948 | 0.308108 | 2.32E-16 |
| S5A_700181210 | 0.252252 | 2.63E-09 |
| S5A_700435349 | 0.097297 | 6.16E-14 |
| S5A_702633797 | 0.223423 | 4.09E-05 |
| S5A_702908675 | 0.227027 | 0.001581 |
| S5A_703364169 | 0.222523 | 0.001969 |
| S5A_704142049 | 0.233333 | 0.003892 |
| S5A_705293051 | 0.100901 | 3.16E-09 |
| S5A_705351694 | 0.102703 | 1.11E-08 |
| S5A_705365208 | 0.097297 | 1.56E-09 |
| S5A_706065949 | 0.276577 | 8.60E-06 |
| S5A_706487909 | 0.407207 | 1.70E-08 |
| S5A_706574241 | 0.384685 | 1.33E-08 |
| S5A_706673002 | 0.338739 | 2.87E-07 |
| S5A_706705101 | 0.285586 | 1.94E-06 |

Supporting information table S6. Location of significant QTL and estimated effects for thousand kernel weight (TKW), estimated test weight, spikelets per spike (SPS), kernel width (K Width), kernel length (K Length), kernel area (K Area) from data collected in the field in 2018 or in the greenhouse in 2017. Data was collected in the field in Raleigh in 2018 for most traits, and in Raleigh and Kinston, North Carolina in 2018 for spikelets per spike.

| Trait | Condition | Chromosome | LOD | Peak Marker | cM | Mean | Effect | SE | % effect |
|---|---|---|---|---|---|---|---|---|---|
| TKW (g) | Field | 4D | 33.76 | RhtD1 | 13 | 27.68 | -2.56 | 0.19 | -9.26 |
| TKW (g) | Field | 3D | 7.4 | S3D_476608044 | 219 | 27.68 | 1.09 | 0.21 | 3.92 |
| TKW (g) | Field | 5A | 6.51 | Awns | 269 | 27.68 | 0.88 | 0.17 | 3.19 |
| TKW (g) | Field | 7A | 4.22 | S7A_71631668 | 82 | 27.68 | 0.79 | 0.19 | 2.85 |
| TKW (g) | Greenhouse | 4D | 23.24 | RhtD1 | 12 | 28.78 | -3.53 | 0.33 | -12.28 |
| TKW (g) | Greenhouse | 2D | 8.1 | PpdD1 | 60 | 28.78 | 1.99 | 0.32 | 6.93 |
| TKW (g) | Greenhouse | 5A | 4.06 | S5A_698528417 | 269 | 28.78 | 1.47 | 0.31 | 5.11 |
| Test Weight (g) | Field | 4D | 33.66 | RhtD1 | 12 | 7.91 | -0.42 | 0.03 | -5.32 |
| Test Weight (g) | Field | 7A | 6.58 | S7A_672150738 | 187 | 7.91 | 0.18 | 0.03 | 2.26 |
| Test Weight (g) | Field | 5A | 5.56 | S5A_698528417 | 269 | 7.91 | 0.15 | 0.03 | 1.89 |
| SPS | Field | 7A | 16.89 | S7A_673996636 | 186 | 20.4 | -0.51 | 0.05 | -2.49 |
| SPS | Field | 2D | 9.11 | PpdD1 | 59 | 20.4 | 0.41 | 0.06 | 2.01 |
| SPS | Field | 5A | 8.44 | S5A_698528417 | 269 | 20.4 | -0.37 | 0.05 | -1.84 |
| SPS | Field | 4D | 6.99 | RhtD1 | 14 | 20.4 | 0.35 | 0.06 | 1.70 |
| SPS | Field | 2B | 5.47 | S2B_600741549 | 108 | 20.4 | 0.24 | 0.05 | 1.15 |
| SPS | Field | 4B | 4.31 | S4B_34025485 | 54 | 20.4 | 0.21 | 0.05 | 1.05 |

| | | | | | | | | | |
|---|---|---|---|---|---|---|---|---|---|
| SPS | Greenhouse | 2D | 25.12 | PpdD1 | 60 | 17.7 | 1.07 | 0.09 | 6.06 |
| SPS | Greenhouse | 4D | 10.6 | RhtD1 | 12 | 17.7 | -0.77 | 0.09 | -4.33 |
| SPS | Greenhouse | 2B | 9.08 | S2B_642534505 | 110 | 17.7 | 0.65 | 0.09 | 3.66 |
| SPS | Greenhouse | 5A | 5.27 | S5A_698528417 | 269 | 17.7 | -0.48 | 0.09 | -2.69 |
| SPS | Greenhouse | 5B | 5.1 | S5B_517797845 | 85 | 17.7 | -0.66 | 0.13 | -3.74 |
| K Width (mm) | Field | 4D | 35.13 | RhtD1 | 12 | 2.98 | -0.09 | 0.01 | -2.94 |
| K Width (mm) | Field | 6A | 5.75 | S6A_115348315 | 69 | 2.98 | 0.04 | 0.01 | 1.22 |
| K Width (mm) | Field | 5A | 5.56 | S5A_73816555 | 58 | 2.98 | 0.03 | 0.01 | 1.17 |
| K Width (mm) | Field | 3D | 4.98 | S3D_496955493 | 229 | 2.98 | 0.03 | 0.01 | 1.02 |
| K Width (mm) | Field | 7A | 4.43 | S7A_71631668 | 81 | 2.98 | 0.03 | 0.01 | 0.86 |
| K Length (mm) | Field | 3D | 13.58 | S3D_298356803 | 206 | 6.28 | 0.1 | 0.01 | 1.56 |
| K Length (mm) | Field | 5A | 9.48 | Awns | 269 | 6.28 | 0.08 | 0.01 | 1.22 |
| K Length (mm) | Field | 1D | 7.7 | S1D_253081308 | 82 | 6.28 | -0.08 | 0.01 | -1.27 |
| K Length (mm) | Field | 3B | 5.67 | S3B_576228566 | 104 | 6.28 | -0.06 | 0.01 | -0.89 |
| K Length (mm) | Field | 6B | 5.01 | S6B_687553875 | 95 | 6.28 | 0.05 | 0.01 | 0.84 |
| K Length (mm) | Field | 1B | 4.72 | S1B_639043667 | 40 | 6.28 | 0.05 | 0.01 | 0.82 |
| K Area ($mm^2$) | Field | 4D | 20.41 | RhtD1 | 14 | 13.97 | -0.55 | 0.05 | -3.90 |
| K Area ($mm^2$) | Field | 3D | 8.73 | S3D_298356803 | 205 | 13.97 | 0.33 | 0.05 | 2.39 |
| K Area ($mm^2$) | Field | 5A | 6.08 | S5A_73816555 | 58 | 13.97 | 0.28 | 0.05 | 2.00 |

| | | | | | | | | | |
|---|---|---|---|---|---|---|---|---|---|
| K Area (mm$^2$) | Field | 6A | 5.34 | S6A_115348315 | 68 | 13.97 | 0.27 | 0.05 | 1.92 |
